# Bioinformatics-based stereochemical elucidation of MM 46115: an unusual antiviral spirotetronate that inhibits clathrin-mediated endocytosis

**DOI:** 10.1101/2025.09.18.677008

**Authors:** Afifah Tasnim, Lauren A. M. Murray, Rebecca A. Clayton, Chidiebere F. Uchechukwu, Jinlian Zhao, Shing C. Liu, Mabilly Cox Holanda de Barros Dias, Ning Liu, Ying Huang, Craig P. Thompson, Lona M. Alkhalaf, Gregory L. Challis, Nicole C. Robb

## Abstract

The influenza virus is a leading cause of respiratory tract infection in humans, with the emergence of antiviral drug resistance posing an ongoing challenge to the treatment of influenza. This underscores the urgent need for effective new antiviral treatments. Spirotetronates are a family of bacterial natural products with a range of potent bioactivities. Some members of this family have antiviral activity. Here, we report the bioinformatics-based stereochemical assignment, antiviral properties, and mechanism of action of the unusual type II spirotetronate MM 46115 (renamed pellemicin). We show that pellemicin is active against both influenza A and B viruses at non-toxic concentrations, and inhibits clathrin-mediated endocytosis, which is required for virus internalisation. Given the need for new treatments for viral infections, the results of our work suggest that pellemicin and related spirotetronates could provide a basis for developing promising alternatives to existing antivirals to combat drug-resistant influenza.

**Graphical abstract:** 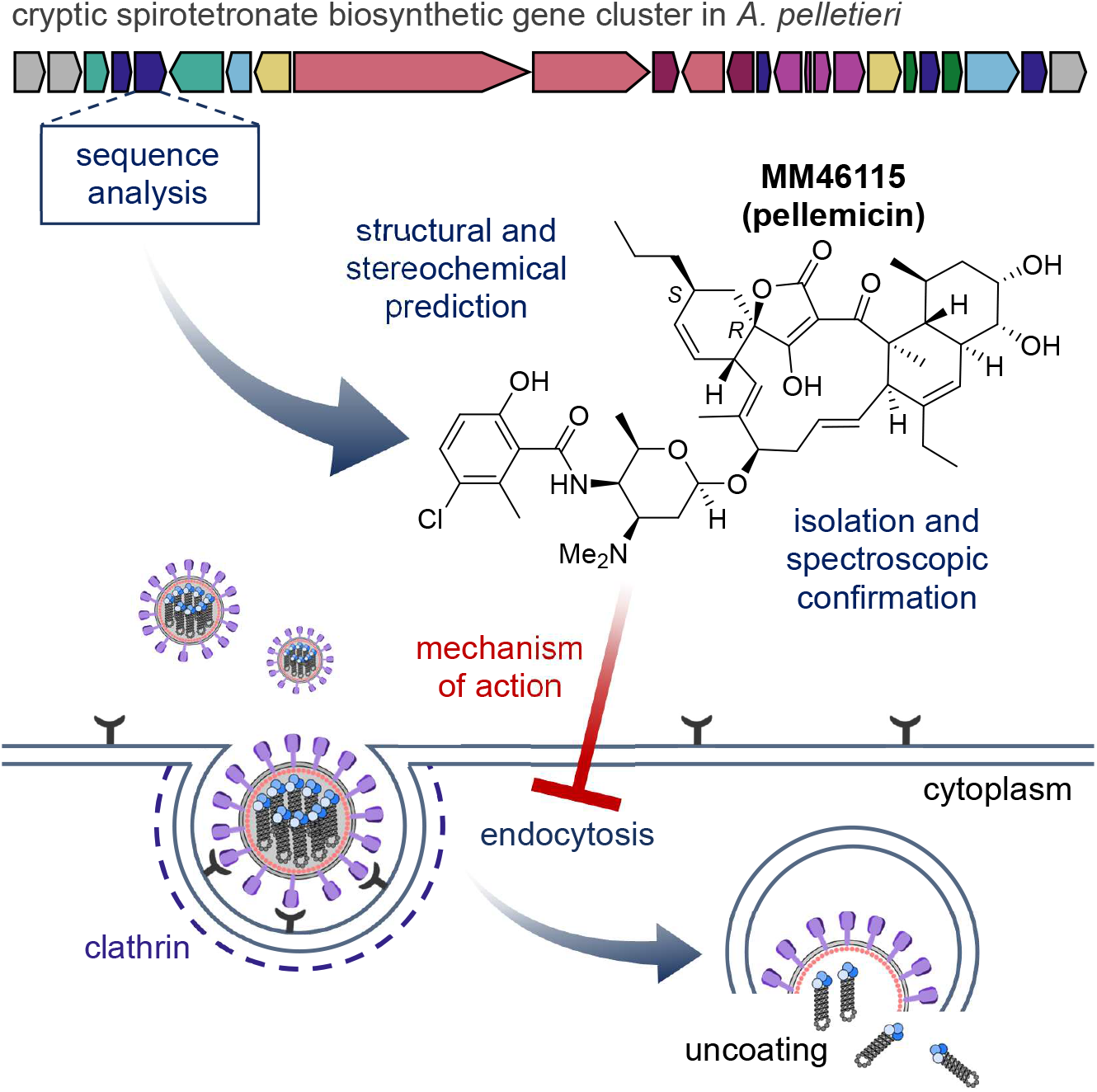

**Highlights:**

- MM 46115 (here renamed pellemicin) produced by a mycetoma pathogen is a promising antiviral
- The pellemicin biosynthetic gene cluster was identified in *Actinomadura pelletieri* DSM 43383
- Sequence analysis of biosynthetic enzymes enabled full stereochemical assignment
- Pellemicin inhibits clathrin-mediated endocytosis, an early stage of the viral life cycle

**Significance:** The influenza virus is a respiratory pathogen with epidemic and pandemic potential, resulting in tens of thousands of deaths each year and causing significant health, social and economic impact. The rise of resistant influenza strains against existing antiviral drugs creates a pressing need for alternative treatments. Natural products - specialised metabolites of bacteria, fungi, and plants that often have antimicrobial activity - are a promising source of therapeutics to treat human disease. Here, we identify a cryptic polyketide biosynthetic gene cluster in the genome sequence of *Actinomadura pelletieri* DSM 43383, an Actinomycete that causes mycetoma in humans. This is proposed to direct production of the previously reported spirotetronate MM 46115 (here renamed pellemicin), which has activity against the influenza virus. Sequence analyses of a polyketide synthase and Diels-Alderase involved in pellemicin biosynthesis, combined with comparative analysis of NMR data for other spirotetronates, enabled us to fully resolve longstanding stereochemical ambiguities in the structure of pellemicin. This underscores the emerging power of bioinformatics-based approaches, and their complementarity to traditional spectroscopic methods, for natural product structure elucidation. We also report extensive insights into the antiviral properties and mechanism of pellemicin, demonstrating that it exhibits antiviral activity at non-toxic doses for up to eight hours after administration and inhibits clathrin-mediated endocytosis, a process that many viruses exploit to enter cells. Only one other bacterial natural product, ikarugamycin, is known to inhibit clathrin-mediated endocytosis. Ikarugamycin contains a tetramate group, which bears a close structural relationship to the tetronate moiety of pellemicin, suggesting that inhibition of clathrin-mediated endocytosis may be a general mechanism of action for antiviral spirotetronates. Our findings therefore indicate that pellemicin, and related spirotetronate natural products, could form a basis for development of new drugs to combat antiviral resistance, and serve as useful tools to study the process of endocytosis.

## Introduction

The influenza virus is a major pathogen in humans and various animal species, with epidemic and global pandemic potential. Influenza infections can result in 650,000 deaths per year in annual epidemics, whilst the Spanish Influenza pandemic in 1918 resulted in 500 million cases and 50 million deaths worldwide, making influenza the biggest viral killer of all time.^1,2^ The main strategy for controlling the spread of influenza is vaccination, but vaccine hesitancy and antigenic drift limits its utility. Patients hospitalised with severe infection are given antivirals. However, the evolution of resistance to these drugs can render them ineffective. For decades, natural product discovery has played a key role in developing new therapeutic agents for a wide range of diseases. The complex structures of these molecules usually endow them with potent biological activities, providing promising starting points for drug development. A recent analysis reveals that, over the last four decades, approximately 40% of newly approved drugs have been either natural products or their derivatives.^3^ Although numerous herbal medicines have been reported to have antiviral activity, natural product remain underexplored as a resource for the discovery of effective new antiviral agents.^4^

Influenza is an enveloped virus with segmented, single-stranded, negative-sense RNA.^5^ Influenza viruses are divided into four main types; A, B, C and D, of which Influenza A and Influenza B are clinically relevant in humans. There are three major types of antiviral active against influenza A that target different parts of the viral lifecycle: M2 channel inhibitors, neuraminidase inhibitors and RNA polymerase inhibitors. M2 channel inhibitors, such as amantadine, block proton translocation, which prevents acidification of the virus core and release of vRNPs from the endosome.^6^ M2 inhibitors are obsolete due to widespread resistance. Neuraminidase inhibitors, such as oseltamivir, block binding of neuraminidase to sialic acid, thus preventing viral release from the cell.^7^ RNA polymerase inhibitors, such as baloxavir marboxil, which are the most recently discovered class of influenza A antivirals, target the viral RdRp complex. Baloxavir acts as a selective inhibitor of the PA subunit of RdRp, preventing the initiation of viral RNA transcription.^8^ Resistance to baloxavir marboxil and oseltamivir results from single amino acid mutations in either the NA protein (H275Y), or the PA protein (PA-I38T).^8,9^ Therefore, novel antivirals with new mechanisms of action are needed to overcome resistance to existing therapies.

Natural products are specialised metabolites produced by plants, bacteria, and fungi. Many of these compounds have evolved to inhibit the growth of competing microorganisms in the environment and, therefore, have useful antimicrobial properties. Spirotetronates are a family of polyketide natural products produced by Actinomycete bacteria that exhibit a range of potent bioactivities.^10^ All known spirotetronates feature an unusual spiro-fused cyclohexene-tetronate embedded in a macrocycle.^10^ In a subset of these metabolites, known as class II spirotetronates, the macrocycle is fused to a functionalised *trans*-decalin. In all spirotetronates studied to date, the spiro-cyclohexene-tetronate moiety results from a Diels-Alder [4+2] cycloaddition between a conjugated *E, E*-diene and an exomethylene tetronate catalysed by a conserved family of β-barrel enzymes. Of the four possible stereochemical outcomes for this reaction (*R*-*endo, S*-*endo, R*-*exo* or *S*-*exo*), only *S*-*exo* and *S*-*endo* have been demonstrated to date for type II spirotetronates (Figure 1A).

**Figure 1.**
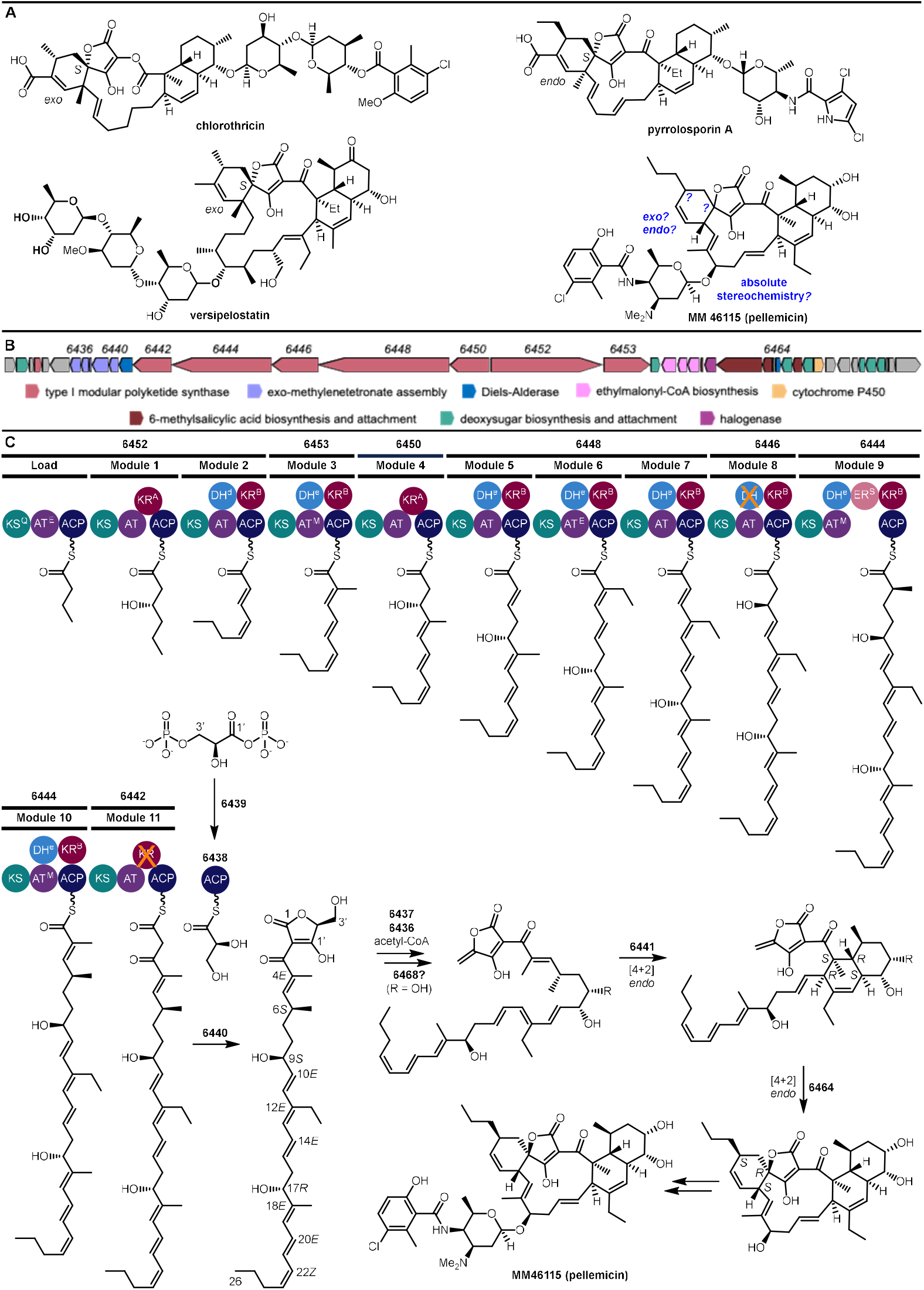
Class II spirotetronates and the proposed biosynthetic pathway to pellemicin. A) Representative members of the class II spirotetronate family of natural products with the stereochemistry of the spirocyclic Diels-Alder adduct highlighted. B) Putative pellemicin biosynthetic gene cluster in *A. pelletieri* DSM 43383. C) Module and domain arrangement of the pellemicin type I modular PKS. The structure of the covalently tethered intermediate predicted to be assembled by each module is shown. D) Post-PKS Diels-Alder reactions catalysed by two successive Diels-Alderases form the pellemicin aglycone.

Numerous spirotetronates, including chlorothricin and pyrrolosporin A (Figure 1A), are active against Gram-positive bacteria.^11,12^ A few have also been reported to possess antiviral activity. These include abyssomicin Y, the quartromicins, 2EPS-A (a C18-methyl congener of pyrrolosporin) and MM 46115.^13,14,15^ Among these, MM 46115, hereafter referred to as pellemicin (Figure 1A), isolated from the opportunistic human pathogen *Actinomadura pelletieri*, was the first type II spirotetronate reported to have antiviral activity.^15^ Although pellemicin was reported to inhibit the growth of influenza A and the respiratory syncytial virus (RSV), cellular cytotoxicity was observed at growth-inhibiting doses.^15^ NMR spectroscopic analysis enabled assignment of the planar structure and the relative stereochemistry of thirteen of the fifteen stereogenic centres in pellemicin.^16^ However, the relative stereochemistry of C2′ and C23, in addition to the absolute stereochemistry, remain unresolved (Figure 1A). Complete stereochemical assignment of pellemicin is necessary to understand what structural features it shares with other type II spirotetronates, enabling initial insights into the relationship between structure and antiviral activity.

Here we report the discovery of a biosynthetic gene cluster (BGC) in *Actinomadura pelletieri* DSM 43383 predicted to assemble a metabolite with a planar structure corresponding to the type II spirotetronate core of pellemicin. High resolution UHPLC-MS analyses of culture extracts, and subsequent purification and NMR spectroscopic analyses, confirmed that *A. pelletieri* DSM 43383 produces pellemicin. Bioinformatics-based predictive stereochemical analyses of a type I modular polyketide synthase and spirotetronate-forming Diels-Alderase encoded by the BGC, combined with comparative ^1^H NMR spectroscopic analysis and circular dichroism (CD) spectroscopy, enabled us to resolve the remaining ambiguities in the structure of pellemicin. Remarkably, these studies indicate that pellemicin is assembled via an intramolecular [4+2] cycloaddition between a (20*E*, 22*Z*)-diene and an exomethylene tetronate, resulting in a (2′*R*, 23*S*)-*endo*-adduct that is extremely rare among known type II spirotetronates. Plaque and cell viability assays showed that pellemicin has potent activity against both influenza A and B viruses over several hours without significant cytotoxicity, and mechanistic studies revealed that it impairs clathrin-mediated endocytosis, an early stage of the viral life cycle. Overall, our findings demonstrate pellemicin is an unusual type II spirotetronate that inhibits influenza virus infection via a rarely observed mechanism among microbial natural products and could provide a useful basis for developing effective new antivirals.

## Results

### Identification and analysis of spirotetronate biosynthetic gene cluster in *A. pelletieri* DSM 43383

During a search of the NCBI database using the BLASTP algorithm, we identified a gene cluster in the draft genome sequence of *A. pelletieri* DSM 43383 (NCBI Accession No. RBWU01000008.1) containing a gene (BZB76_6464) encoding a protein with similarity to Diels-Alderases known to catalyse spirocycle formation in the biosynthesis of several spirotetronates (Figure 1B). Comparative sequence analysis of the proteins encoded by other genes in the BGC enabled us to predict generalised functions for most of them (Figure 1B and Table S1). Seven genes (BZB76_6442 to BZB76_6453) encode type I modular polyketide synthase (PKS) subunits, while a five-gene cassette (BZB76_6436 to BZB76_6440) encodes homologues of proteins known to create and fuse a *D*-glyceryl thioester to β-keto thioester products of modular PKSs resulting in formation of a 2-acyl-4-exomethylenetetronic acid (Figure 1B). The BGC also contains genes encoding a halogenase and a cytochrome P450 (CYP), and enzymes for the biosynthesis of (2*S*)-ethylmalonyl-CoA, a 6-methylsalicyl moiety, and a 3,4-diamino-3,4,6-trideoxy-α-*D*-glucose derivative (Figure 1B). In addition to the gene encoding the spirocycle-forming Diels-Alderase the BGC contains a gene (BZB76_6441) encoding a decalin-forming Diels-Alderase involved in type II spirotetronate biosynthesis (Figure 1B).

The PKS/NRPS Analysis server (https://nrps.igs.umaryland.edu/) was used to predict the module and domain organisation of the type I modular PKS encoded by BZB76_6442 to BZB76_6453.^17^ The sequences of each catalytic domain type were aligned to provide detailed insights into their function / substrate specificity / stereospecificity (Figure S1). The presence of a ketosynthase domain with the active site Cys residue mutated to Gln (KS^Q^) in the first module of the PKS subunit encoded by BZB76_6452 indicates this is the loading module, which initiates polyketide chain assembly by catalysing decarboxylation of the (alkyl)malonyl thioester attached to the acyl carrier protein (ACP) domain, resulting in formation of the starter unit.^18^ This subunit is also predicted to contain chain extension modules 1 and 2 (Figure 1C). All chain extension modules are predicted to contain functional ketoreductase (KR) domains, except the module encoded by BZB76_6442 (Figure S1). The KR domain in this module contains several deletions and lacks the catalytically important active site Tyr residue (Figure S1). Thus, the KR domain is predicted to be inactive and BZB76_6442 is proposed to encode the eleventh and last chain extension module, because it is the only one capable of producing the β-keto thioester required for chain release via tetronate formation (Figure 1C).^19^ The order of the other subunits in the PKS is proposed to reflect the organization of the genes encoding them; BZB76_6453, which is directly downstream of BZB76_6453 encodes the second subunit (containing chain extension module 3), and BZB76_6450-6444, which are directly upstream of BZB76_6442 encode the third, fourth, fifth and sixth subunits (containing chain extension modules 4, 5-7, 8, and 9-10, respectively) (Figure 1C).

A sequence alignment predicts that the acyltransferase (AT) domains in the loading module and chain extension module 6 are specific for (2*S*)-ethylmalonyl-CoA, whereas the AT domains in chain extension modules 3, 9 and 10 are specific for (2*S*)-methylmalonyl-CoA and those in the remaining modules utilize malonyl-CoA (Figure S1).^20^ Thus, the acyl chain built by the PKS is hypothesized to derive from a butanoyl starter unit, and contain methyl branches at C4, C6 and C18 and an ethyl branch at C12 (Figure 1C). Similarly, a sequence alignment predicts the KR domains in chain extension modules 1 and 4 are “A1-type”, while those in the other chain extension modules are “B1-type” (Figure S1).^21^ All the modules containing a “B1-type” KR domain also contain a functional dehydratase (DH) domain, except chain extension module 8, in which the universally conserved active site Asp residue has been mutated to Glu (Figure S1). The DH domain-containing modules, with the exception of module 2, are thus all expected to install an *E*-configured enoyl thioester.^22^ The YGP motif in canonical enoyl thioester-forming domains, is mutated to WGP in the module 2 DH domain (Figure S1), indicating that it forms a 2, 4-dienoyl thioester.^23^ The substrate for this diene-forming DH domain is predicted, on the basis of the KR domain stereospecificity assignments, to be the (3*R*, 5*S*)-3, 5-dihydroxy thioester. Thus, the (2*E*, 4*Z*)-2, 4-dioenyl thioester is proposed to be assembled by chain extension module 2 (Figures 1C and S2). Sequence analysis of the enoylreductase (ER) domain in module 9 indicates it catalyses formation of a 2*S*-configured 2-methyl thioester (Figure S1).^24^ Overall, these analyses predict that the PKS assembles a (4*E*, 6*S*, 9*S*, 10*E*, 12*E*, 14*E*, 17*R*, 18*E*, 20*E*, 22*Z*)-β-keto thioester, which undergoes condensation with a glyceryl-ACP thioester to give the corresponding 2′-hydroxymethyltetronic acid (Figure 1C). Conversion of this intermediate to the 2′-exomethylenetetronic acid, via established enzymology,^25^ and a [4+2] cycloaddition analogous to that reported for other type II spirotetronates and tetramates is predicted to result in decalin formation via an *endo* transition state (Figure 1C).^26^

The absolute configurations of all the stereocentres in the proposed decalin intermediate match the relative configurations assigned, based on NMR spectroscopic analyses,^16^ to the corresponding stereocentres in pellemicin (with the exception of the C8 stereocentre in pellemicin, which results from a CYP-catalysed hydroxylation reaction – neither the timing nor the stereoselectivity of this reaction can be predicted). Thus, we proposed that the cryptic type II spirotetronate BGC we identified in the *A. pelletieri* DSM 43383 draft genome sequence directs the biosynthesis of pellemicin, which has 4*S*, 5*R*, 6*S*, 8*R*, 9*S*, 12*R*, 17*R* absolute configuration. The relative stereochemistry of C23 and C2′ in pellemicin, could not be assigned during the previous NMR spectroscopic study due to a lack of diagnostic signals in the nuclear Overhauser effect spectrum.^16^ The stereochemistry of C20, C23 and C3′ depends on (i) the configuration of the C20-C21 and C22-C23 double bonds, which we assigned in our predictive analysis of the PKS (see above), and (ii) the stereochemical course of the spirotetronate-forming [4+2] cycloaddition, which can proceed via either an *exo* or *endo* transition state involving addition of the diene to either the *re* or *si* face of the exomethylene double bond in the tetronate (the corresponding products are designated *R*-*exo, R*-*endo, S*-*exo* and *S*-*endo*, respectively). To address the question of the unresolved C23 and C2′ stereochemistry, we conducted a phylogenetic comparison of the Diels-Alderase encoded by BZB76_6464 with those encoded by BGCs for other type II spirotetronates with *R*-*endo, S*-*exo* and *S*-*endo* configurations (Figure 2A). Based on this, we hypothesised that the Diels-Alderase encoded by BZB76_6464 catalyses formation of an *R*-*endo* adduct with 2′*R*, 20*S*, 23*S* configuration (Figure 1C).

**Figure 2:**
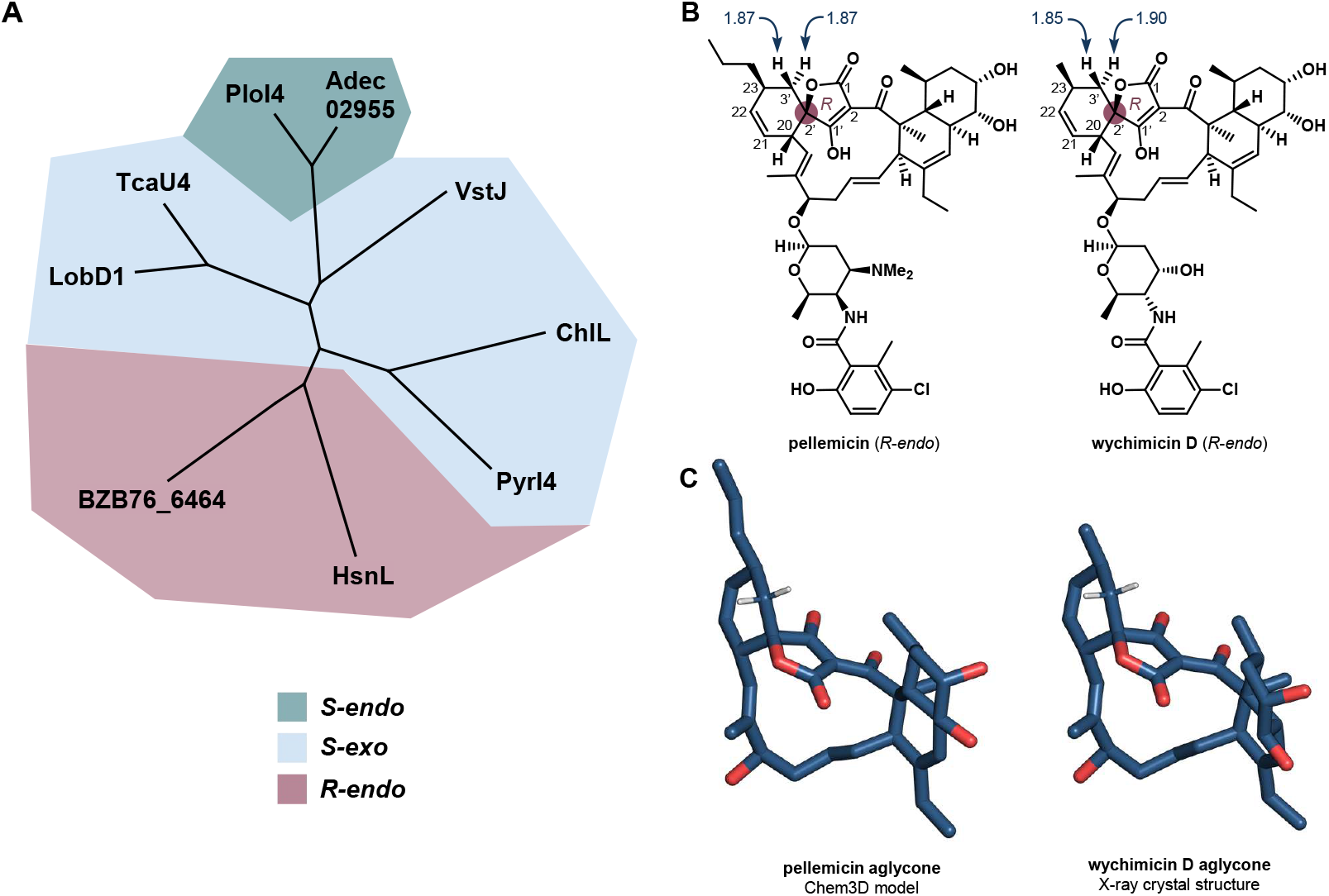
Comparison of chemical shifts for C3 protons in pellemicin and wychimicin D. A) Phylogenetic comparison of the sequences of Diels-Alderases catalysing the formation of spirotetronates with different stereochemistry in type II spirotetronate biosynthesis. Details of the compound assembled are given in Table S5. B) Comparison of the relative and absolute stereochemistry proposed for pellemicin with that elucidated by X-ray crystallography for wychimicin D. The signals at δ_H_ = 1.85/1.90 and 1.87/1.87 ppm for the diastereotopic C3′ protons in wichimycin D and pellemicin, respectively, which result from the 2′*R*, 20*S*, 23*S* stereochemistry, are highlighted. C) Comparison of a molecular mechanics model of the pellemicin aglycone with the aglycone portion of the wychimicin D X-ray crystal structure.

### *A. pelletieri* DSM 43383 produces pellemicin enabling experimental confirmation of predicted stereochemistry

To confirm that the cryptic spirotetronate BGC identified in the draft genome sequence of *A. pelletieri* DSM 43383 directs pellemicin biosynthesis, the strain was grown for 10 days at 28 °C on GYM (glucose, yeast and malt extract) agar medium, and the culture was extracted with ethyl acetate and analysed using UHPLC-Q-ToF-MS (Figure S3). Ions with *m/z* = 891.4557 and 913.4376 were observed, corresponding to the pellemicin [M+H]^+^ and [M+Na]^+^ ions (Figure S3). The corresponding metabolite was purified from large scale culture extracts using reverse phase HPLC and ^1^H, ^13^C (JMOD), COSY, HSQC, HMBC, NOESY, ROESY and TOCSY NMR spectra were recorded (Figures S4-S11). The δ_H_ and δ_C_ values for the purified metabolite were in excellent agreement with those previously reported for pellemicin (Table S2).^16^ Identical chemical shifts were observed for the diastereotopic protons attached to C-3′. This prevented stereochemical assignment of C23 and C2′ relative to C20 using a combination of NOESY/ROESY and coupling constant data.

We thus turned our attention to comparison of ^1^H NMR spectroscopic data for pellemicin with related metabolites, formed via stereodivergent spirotetronate-forming [4+2] cycloadditions, that have been stereochemically elucidated using X-ray crystallography.^27,28,29^ The diastereotopic C3′ protons in the *S-exo* spirotetronate glenthmycin B and the *S-endo* spirotetronate pyrrolosporin A have chemical shift differences of 0.44 and 0.78 ppm, respectively (Figure S12). Both the C2′–O and C23-C24 bonds are *anti* to one of the C3′–H bonds and *gauche* to the other in the X-ray crystal structure of the glenthmycin B aglcycone (Figure S12). Similarly, in the X-ray crystal structure of pyrrolosporin A, the C23-C24 bond is also oriented *anti* to one of the C3′–H bonds and *gauche* to the other, whereas the C2′–O bond bisects the C3′–H bonds, lying gauche to both (Figure S12). The differences in chemical shift values for the C3′ protons in glenthmycin B and pyrrolosporin A can be attributed to the asymmetric distribution of one or both vicinal substituents. In contrast, both the C2′–O and C23-C24 bond bisect the C3′–H bonds in the X-ray crystal structure of the *R-endo* spirotetronate wychimicin D (Figure 2B). Thus, both protons attached to C–3′ experience similar deshielding / shielding effects from the neighbouring C–O and C-C bonds, as evidenced by their almost identical chemical shift values (Figure 2B).

A molecular mechanics model of the pellemicin aglycone structure predicted using bioinformatics (see above) adopts a very similar conformation to the X-ray crystal structure of the wychimicin D aglycone (Figure 2C). Consequently, the C2′–O and C23-C24 bonds both bisect the C3′–H bonds in the model, consistent with the diastereotopic C3′ protons in pellemicin having identical chemical shift values (Figure 2C).The planar structures of pellemicin and wychimicin D aglycones differ only in the C23 substituent, which is an *n*-propyl group in the former and a methyl group in the latter. Virtually all protons and carbons in the pellemicin and wychimicin aglycones have very similar chemical shift values (Table S3). C22, C23, and C24 are the only carbons with signals differing by >1 ppm in ^13^C NMR spectrum, as expected for the substitution of an *n*-propyl group with a methyl group at C23. Overall, the spectroscopic data are fully consistent with our bioinformatics-based assignment of the 2′*R*, 20*S*, 23*S* relative and absolute configuration to pellemicin (Figure 2C). To further confirm our absolute stereochemical assignment, we compared the measured Electronic Circular Dichroism (ECD) spectrum of pellemicin to the calculated spectrum of its aglycone (Figure S13 and Table S4).

### Pellemicin inhibits influenza-induced cell death

Having established the complete relative and absolute stereochemistry of pellemicin, we turned our attention to the effect of this compound on the growth of the influenza A virus. A stock solution of pellemicin was produced by dissolving the compound in dimethyl sulfoxide (DMSO) and dilutions of the stock, ranging from 0.001235 - 123.5 µg, were made in cell culture media, before being mixed with influenza A/WSN/33 and added to MDCK cells. An agarose overlay was added to the cells and plaques were stained and counted after three days. Addition of A/WSN/33 virus to the MDCK cells in the absence of pellemicin resulted in a high number of plaques, whilst a negative control lacking virus produced no visible plaques (Figure 3A). Addition of increasing concentrations of pellemicin to the infected MDCK cells resulted in a decrease in the number of plaques formed (Figure 3A, upper panel), whereas no plaques were visible when the compound was added to non-infected cells (Figure 3A, lower panel), confirming that pellemicin is not contributing to plaque formation or causing cell death. DMSO alone was not found to influence virus titres (Figure S16). The virus titre was quantified in each condition and found that addition of lower concentrations of pellemicin (0.01235 µg and 0.001235 µg) resulted in a small, but non-significant, reduction in viral titres, suggesting that these concentrations are too dilute to cause robust viral inhibition (Figure 3B). However, a range between 12.35 and 0.1235 µg pellemicin produced significantly fewer plaques and lower titres compared to untreated cells infected with virus (Figure 3B). Overall, this indicates that pellemicin results in a dose-dependent reduction in A/WSN/33 virus titre.

**Figure 3.**
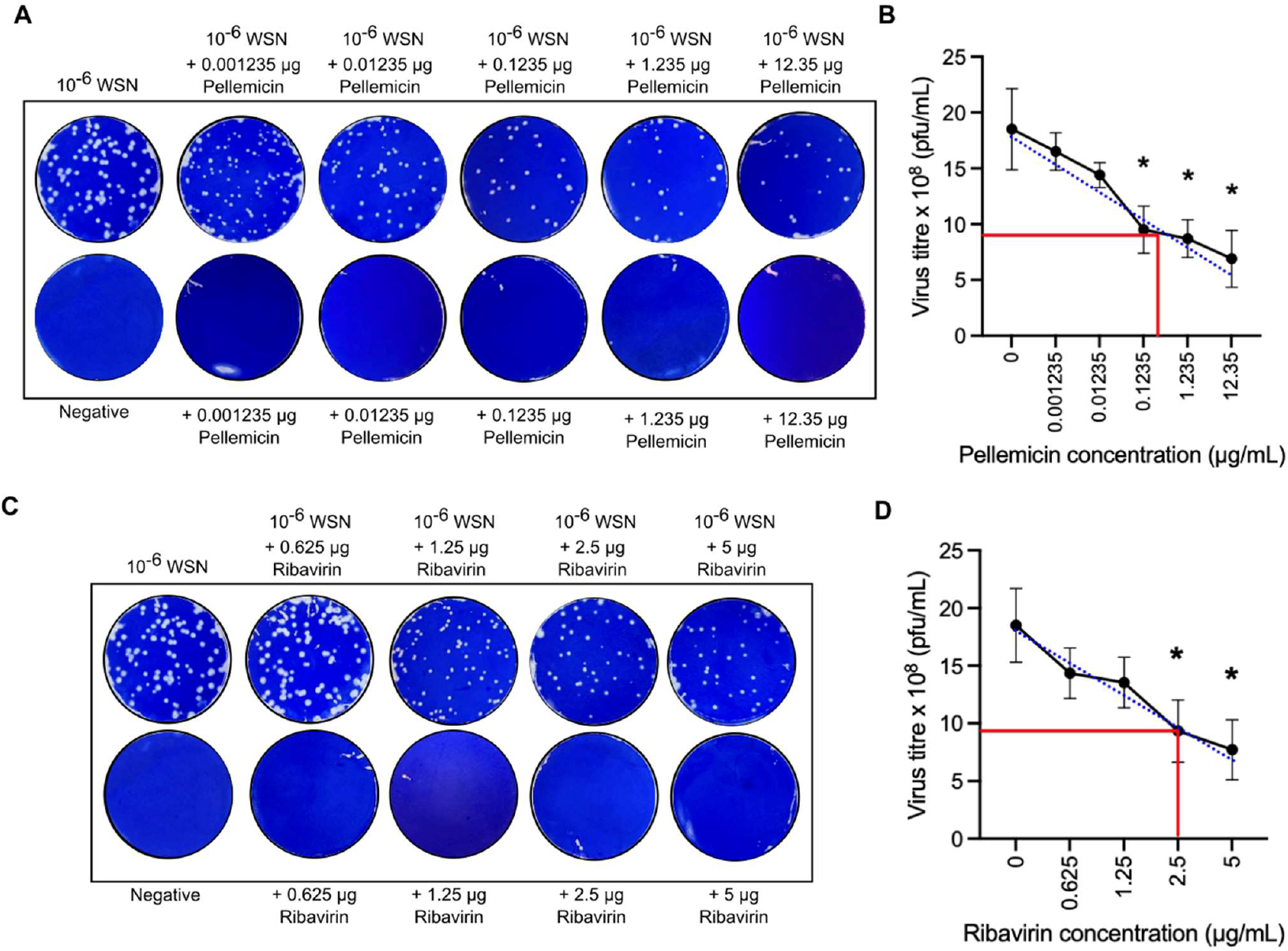
Pellemicin inhibits influenza induced cell death. A) MDCK cells were infected with influenza A virus A/WSN/33 at an MOI of 0.1 (top) or cultured in DMEM/0.5% FBS alone (bottom) and treated with pellemicin (0.001235 - 12.35 µg/mL) or left untreated before being subjected to plaque assay. Representative images of plaque assays are shown. B) Mean virus titre of infected cells in the presence of increasing concentrations of pellemicin, from three experiments. Error bars show standard deviation, blue is line of best fit. Titres were compared to the untreated group using a one-way ANOVA, * p<0.05 was considered statistically significant. The IC_50_ value was calculated at 0.3396 µg for pellemicin (red line). C) MDCK cells were infected with influenza A virus A/WSN/33 at an MOI of 0.1 (top) or cultured in DMEM/0.5% FBS alone (bottom) and treated with ribavirin (0.625 -5 µg/mL) or left untreated before being subjected to plaque assay. Representative images of plaque assays are shown. D) Same as B) but for ribavirin treated cells. The IC_50_ value was calculated at 2.5 µg for ribavirin (red line).

The activity of pellemicin against influenza A was compared with ribavirin, a broad-spectrum antiviral which prevents viral mRNA synthesis and has previously been shown to be effective against the influenza A virus.^30^ Addition of increasing concentrations of ribavirin to the infected MDCK cells resulted in a decrease in the number of plaques formed (Figure 3C, upper panel), whilst no plaques were visible when the compound was added to non-infected cells (Figure 3C, lower panel). The virus titre was quantified for each condition and found that 2.5 and 5 µg ribavirin produced significantly fewer plaques and lower titres compared to the untreated control, thereby demonstrating antiviral activity at these concentrations (Figure 3D). The IC_50_ value (concentration of inhibitor at which number of plaques is reduced by 50%) for pellemicin was 0.3396 µg (0.761 µM) whilst for ribavirin this was 2.5 µg (10.2 µM) (Figure 3B&D), suggesting that the potency of pellemicin against influenza A is 13-fold higher than ribavirin.

### Pellemicin demonstrates antiviral activity at non-toxic concentrations

To establish the cytotoxic properties of pellemicin on healthy cells we undertook a neutral red cell cytotoxicity assay to measure cell viability. Neutral red is a eurhodin dye which stains lysosomes in viable cells, whereas non-viable cells cannot take up the chromophore. Decreasing concentrations of pellemicin (123.5 – 0.01235 μg) or ribavirin (5 – 0.625 μg) were added in triplicate to confluent A549 cells, a human lung epithelial cell line. After washing, the amount of released dye from viable cells in each condition was used to determine the total number of viable cells, or drug cytotoxicity. Both the highest concentration conditions of pellemicin and ribavirin were found to significantly reduced cell viability by approximately 20% compared to the negative media-only control, however lower concentrations of both compounds did not cause significant cell death (Figure 4A). To further validate these results, a known cytotoxic drug, doxorubicin, and the A/WSN/33 virus (1 x 10^6^ PFU) were both shown to cause significant cell death, as expected. Similar results were also obtained using MDCK cells, a canine kidney cell line (Figure S17).

**Figure 4.**
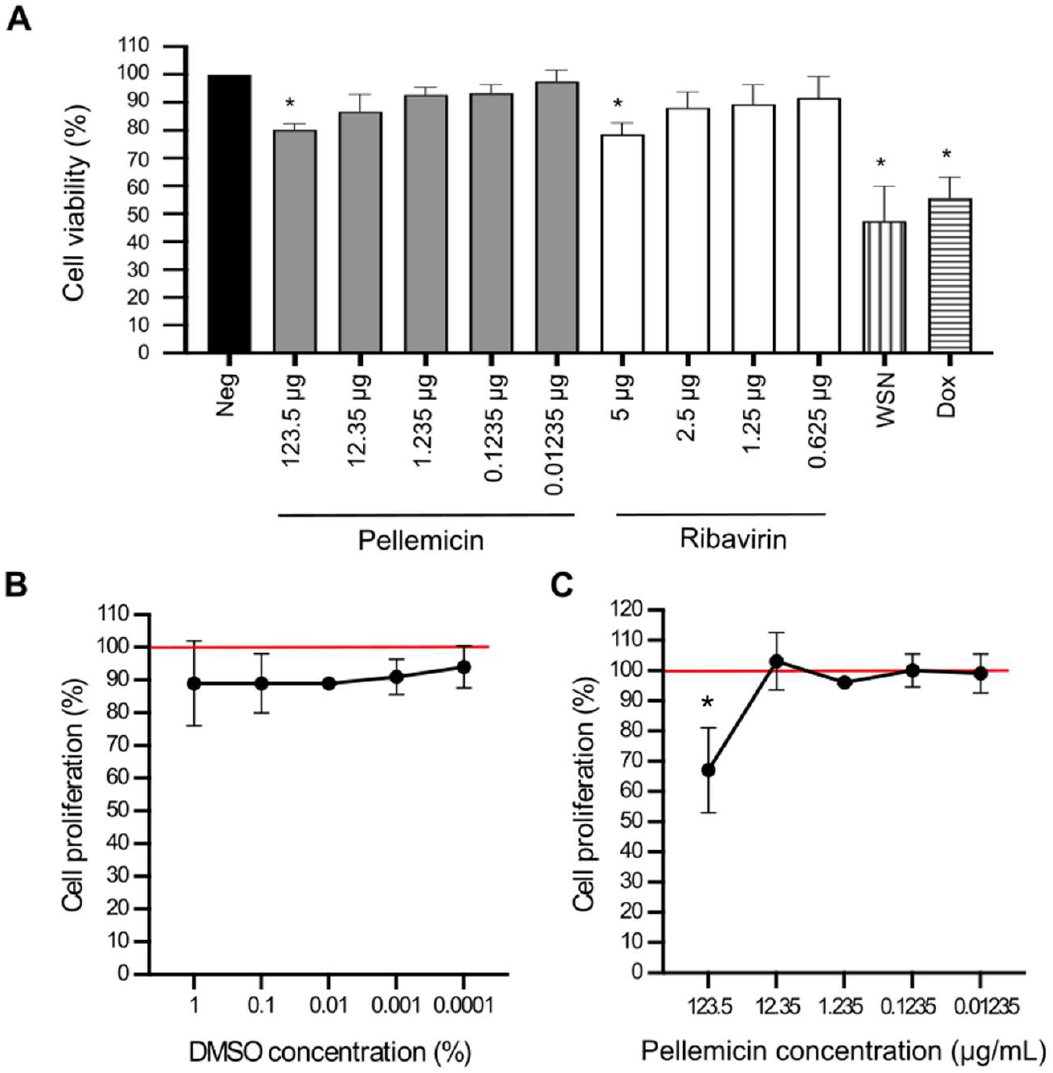
Cytotoxicity of pellemicin measured by cell viability and cell proliferation assays. A) A549 cells were left untreated (negative control), treated with pellemicin (123.5 – 0.01235 µg/mL), ribavirin (5 – 0.625 µg/mL), infected with A/WSN/33 (WSN), or treated with doxorubicin (Dox; 0.04 mM). Cell viability was measured using a neutral red assay kit in triplicate and absorbance measured at 540 nm. B) MDCK cells were treated with DMSO (1-0.0001 %) or left untreated (negative, red line). A cell proliferation assay based on the measurement of thymidine analogue BrdU incorporation during DNA synthesis was carried out in triplicate and absorbance measured at 450 nm. C) Same as B), however MDCK cells were treated with pellemicin (123.5 – 0.01235 µg/mL) or left untreated (negative, red line). The negative control was set at 100% in both assays and groups compared with a one-way ANOVA (*p<0.05). Error bars show standard deviation

To verify our results with a second method, we used a colorimetric cell proliferation assay based on the measurement of thymidine analogue 5-bromo-2′-deoxyuridine (BrdU) incorporation during DNA synthesis in replicating cells. Decreasing concentrations of pellemicin (123.5 – 0.01235 μg) diluted in DMSO, or DMSO only, were incubated with MDCK cells before an anti-BrdU antibody was used to quantify the degree of BrdU incorporation. We started by measuring the effect of decreasing concentrations of DMSO (1 – 0.0001%) only and found that even high concentrations of DMSO did not significantly affect cell proliferation (Figure 4B). Next, we found that only the highest concentration of pellemicin (123.5 µg) reduced cell proliferation by 33% compared to the negative, whereas no changes were seen with lower concentrations of the compound (Figure 4C). Taken together, data from both our cell viability and cell proliferation assays showed that 0.1235 µg pellemicin did not significantly reduce cell viability, indicating that pellemicin can produce antiviral activity without cytotoxicity.

### Pellemicin inhibits influenza for up to eight hours post-infection

To assess the duration of pellemicin antiviral activity, MDCK cells were infected with either influenza A (A/WSN/33) or influenza B (B/Beijing/1/87) at a multiplicity of infection (M.O.I.) 0.01, in the presence or absence of the compound. The cell supernatants were harvested at 2-, 4-, 6-, 8-, 12- and 24-hours post-infection and titred via plaque assay. For influenza A, the 2-, 4- and 6-hour infections carried out with pellemicin present produced fewer plaques compared to the same timepoints without pellemicin (Figure 5A). However, the 8-, 12- and 24-hour infections produced a visually similar number of plaques in the presence or absence of pellemicin. When a second dose of pellemicin was added to the infected cells at 6-hours post-infection the reduction in plaque numbers was maintained for longer, suggesting that the restoration of viral titre at later timepoints is due to the metabolism or breakdown of the compound after this time. Pellemicin also showed good activity against an influenza B virus, with supernatants harvested at 2-, 4-, 6- and 8-hour post-infection in the presence of pellemicin producing fewer plaques compared to the same timepoints without pellemicin (Figure 5B). Quantification of the viral titres confirmed that pellemicin treatment inhibits influenza A-induced cell death for up to 6 hours and influenza B-induced cell death for up to 8 hours in MDCK cells (Figure 5C & D).

**Figure 5.**
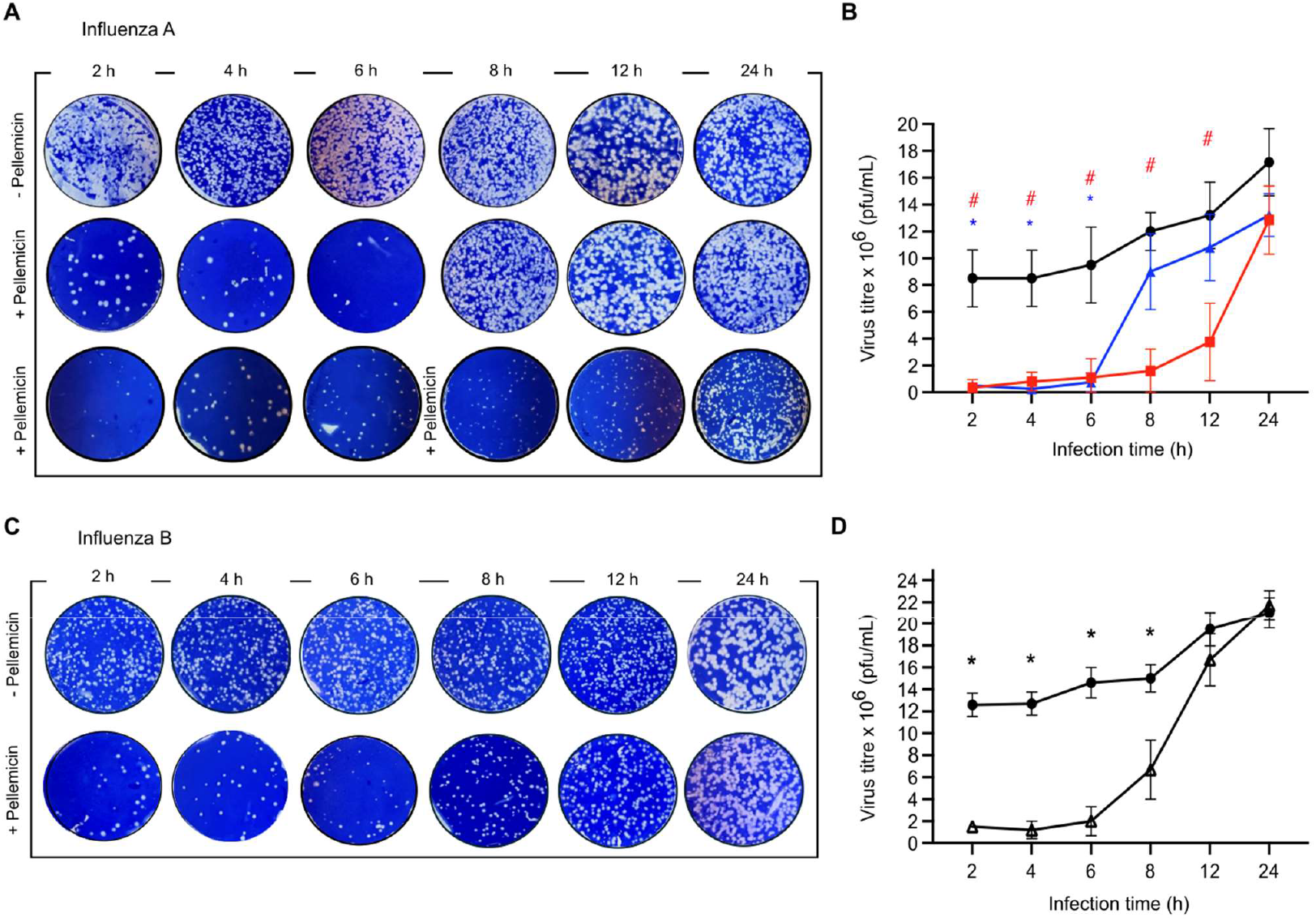
Pellemicin inhibits influenza replication for up to 8 hours. A) MDCK cells were infected with influenza A/WSN/33 virus (MOI 0.1) and treated with 0.1235 µg/mL pellemicin once at 0 hours, twice at 0 and 6 hours or left untreated for 2, 4, 6, 8, 12 and 24 hours. The cell supernatant was harvested at each timepoint until 24 hours and titred via plaque assay. B) Mean virus titres of the treatment groups (one treatment - blue triangles (* p<0.05), two treatments - red squares (# p<0.05)) were compared at each timepoint to the untreated group (black circles) using an unpaired t-test. Error bars show standard deviation from three independent repeats. C&D) Same as A) and B) except for the influenza B/Beijing/1/87 virus (MOI 0.1).

### Pellemicin targets an early stage of the influenza virus life cycle

Next, we carried out a series of experiments to investigate the stage of the virus life cycle at which pellemicin exerted its antiviral activity. Confluent MDCK cells were infected with influenza A/WSN/33, and either left untreated or treated with pellemicin either at the time of infection (0 hours) or at 2- or 4-hours post-infection. The viral supernatants were harvested and titred by plaque assay. Treating cells with pellemicin at the time of infection reduced plaque formation and virus titre compared to untreated cells, however, adding pellemicin post-infection did not (Figure 6A & B). To test whether addition of pellemicin prior to infection resulted in antiviral activity, cells were treated for 2 hours, followed by a wash step, before virus was added. Preincubation of the cells with pellemicin before infection reduced plaque formation and viral titres to a similar extent to simultaneous addition of pellemicin and virus (Figure 6C & D). This suggests that pellemicin does not need to interact with the virus prior to infection to cause inhibition.

**Figure 6.**
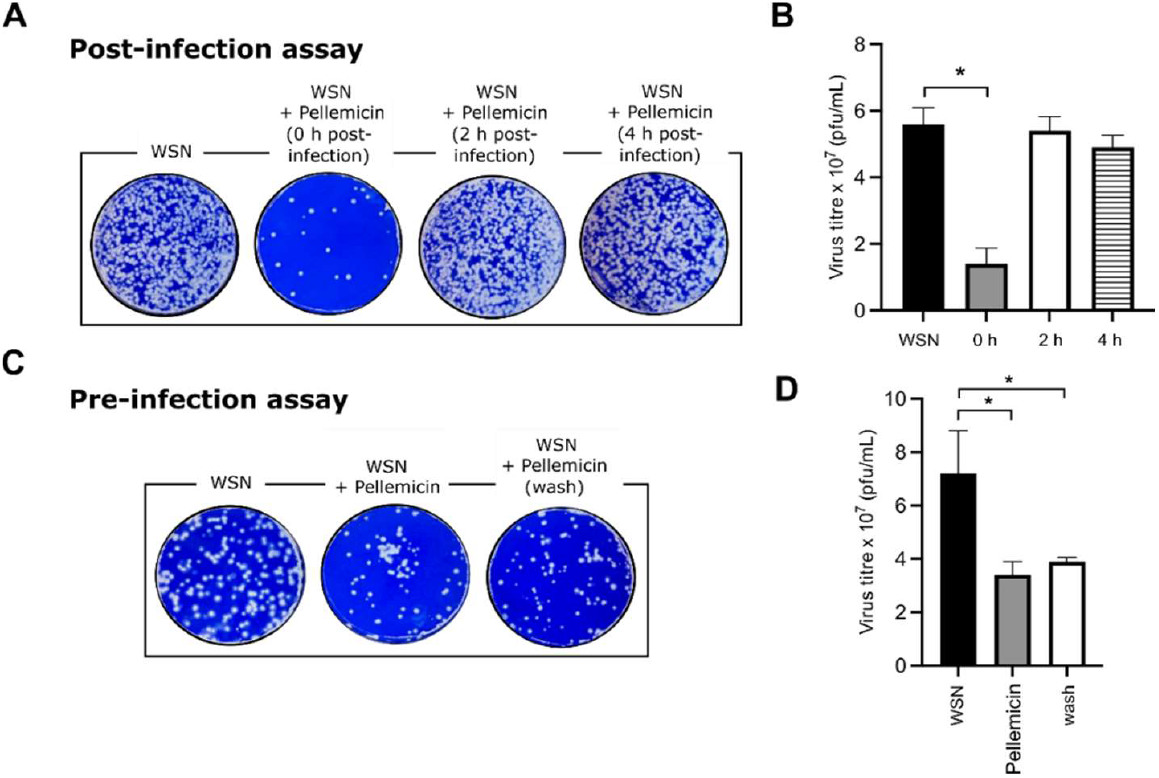
Pellemicin is effective when added before or during infection. A) Effect of pellemicin treatment during or post-infection. pellemicin (0.1235 µg/mL) was added to MDCK cells infected with WSN virus (MOI 0.01) at 0-, 2- and 4-hours post-infection. Cells infected with virus only were included as an untreated control. Infections were performed for 6 hours in total before the supernatant was harvested and titred via plaque assay. B) Mean virus titres of treatment groups compared to untreated control using an unpaired t-test. * p<0.05 was considered statistically significant. Error bars show standard deviation from three independent repeats. C) Effect of pre-incubation of cells with pellemicin. Preincubation of cells with pellemicin for 2 h, with and without washing with PBS, before infection with A/WSN/33 virus. Cells infected with WSN alone were used as an untreated control. Infections were performed for 6 hours in total before the supernatant was harvested and titred via plaque assay. D) Mean virus titres of treatment groups compared to untreated control using an unpaired t-test. * p<0.05 was considered statistically significant. Error bars show standard deviation from three independent repeats.

This result was further supported by a haemagglutination assay. When red blood cells (RBCs) are added to V-bottom plates they settle to form clots. Addition of virus to the RBCs resulted in haemagglutination -the binding of the virus haemagglutinin (HA) to *N*-acetylneuraminic acid (a type of sialic acid) on the RBC surface, causing the RBCs to cross-link and thus preventing clotting (Table 1A, Figure S18A). Addition of increasing concentrations of pellemicin to the virus stock still resulted in haemagglutination, even at the highest inhibitor concentration (Table 1B, Figure S18B). This suggests that pellemicin is not inhibiting haemagglutination, and therefore not preventing the viral HA protein from binding to the RBCs. Addition of compound alone still allowed the RBCs to form clots, suggesting that pellemicin does not interact with sialic acid on the RBC surface either (Table 1C, Figure S18C). Furthermore, DMSO only did not affect the RBCs, which rules out the possibility of DMSO causing RBC lysis (Table 1D, Figure S18D). A positive control with serum from a patient immunised with the influenza vaccine was used as a control. This serum contains anti-HA antibodies which can bind to the viral HA and block it from interacting with the RBCs, resulting in clotting (Table 1E & F, Figure S18E & F). We therefore concluded that although pellemicin works at an early stage of the viral life cycle it does not block the binding of viral HA to host sialic acid to cause antiviral activity.

**Table 1.** Effect of pellemicin on agglutination of red blood cells. A/WSN/33 was mixed with red blood cells in the presence or absence of pellemicin (A&B). Control groups with pellemicin alone (C) or DMSO (D) were included. A positive control with serum from a patient immunised with the influenza vaccine was included, with virus (E) and without (F). Haemagglutination activity was tested by visually assessing haemagglutination. Results indicate haemagglutination (H), haemagglutination inhibition (HI) or no reaction (NR).

| **A** |  | **B** |  |
| --- | --- | --- | --- |
| **Virus alone** | **Result** | **Virus stock + pellemicin** | **Result** |
| Stock | H | Virus | H |
| 10<sup>-1</sup> | NR | Virus + 123.5 µg | H |
| 10<sup>-2</sup> | NR | Virus + 12.35 µg | H |
| 10<sup>-3</sup> | NR | Virus + 1.235 µg | H |
| **C** |  | **D** |  |
| **Pellemicin alone** | **Result** | **DMSO alone** | **Result** |
| 123.5 µg | NR | 1% | NR |
| 12.35 µg | NR | 0.1% | NR |
| 1.235 µg | NR | 0.01% | NR |
| 0.1235 µg | NR | 0.001% | NR |
| **E** |  | **F** |  |
| **Virus + serum** | **Result** | **Serum alone** | **Result** |
| Virus | H | Serum stock | NR |
| Virus + serum stock | HI | Serum (1:2) | NR |
| Virus + serum (1:2) | HI | Serum (1:4) | NR |
| Virus + serum (1:4) | HI |  |  |

To support this further, we investigated the effect of pellemicin addition on later stages of the viral lifecycle. First, qRT-PCR was used to measure the levels of intracellular viral RNA in pellemicin-treated and untreated cells over 24 hours. RNA extraction was performed 2-24 hours after viral infection and RNA levels in supernatants and cell lysates were measured. No significant differences between RNA levels in treated and untreated groups were observed over 24 hours, in either cell lysates (Figure S19A) or supernatants (Figure S19B). Next, a neuraminidase (NA) inhibition assay was used to examine whether pellemicin targets the viral NA protein. Addition of oseltamivir, a licenced NA inhibitor, is known to bind to the virus NA protein and prevent it from catalysing the hydrolysis of the fluorogenic MUNANA substrate into the fluorescent 4-methylumbelliferone product, therefore resulting in a dose-dependent reduction in fluorescence levels (Figure S20A). We calculated an IC_50_ value of 10 nM for oseltamivir, which supports findings in other studies.^31^ However, addition of pellemicin did not cause a dose-dependent reduction in fluorescence levels, which suggests the viral NA is still active in the presence of pellemicin, and the substrate is still being hydrolysed into the fluorescent product (Figure S20B). Together our data suggests that pellemicin does not affect viral replication or NA activity, which are later stages of the virus cycle.

### Pellemicin inhibits clathrin-mediated endocytosis

Clathrin-mediated endocytosis is the most common endocytic pathway and a major internalization pathway for influenza virus entry into host cells, however filamentous strains of influenza have been shown to utilise macropinocytosis as an alternate entry pathway.^32,33^ To assess whether endocytosis was affected by pellemicin addition we tested the effect of compound addition on A/WSN/33, a spherical influenza strain, and A/Udorn/72, a well characterised influenza strain exhibiting both a spherical and filamentous morphology (Fig 7A).^34^ MDCK cells were infected with either A/WSN/33 or A/Udorn/72, in the presence or absence of pellemicin, before the cell supernatants were harvested and titred via plaque assay (Fig 7B). A/Udorn/72 exhibited significantly smaller plaques than A/WSN/33, a distinguishing feature of this virus (Fig 7B). For A/WSN/33 the infections carried out with pellemicin present produced ∼ 50% fewer plaques compared to infection without pellemicin as expected, however, infection with A/Udorn/72 produced a similar number of plaques in the presence or absence of compound (Figure 7C). This suggests that a viral strain that uses endocytosis to enter cells is affected by pellemicin addition, whilst a viral strain that utilises macropinocytosis is not.

**Figure 7.**
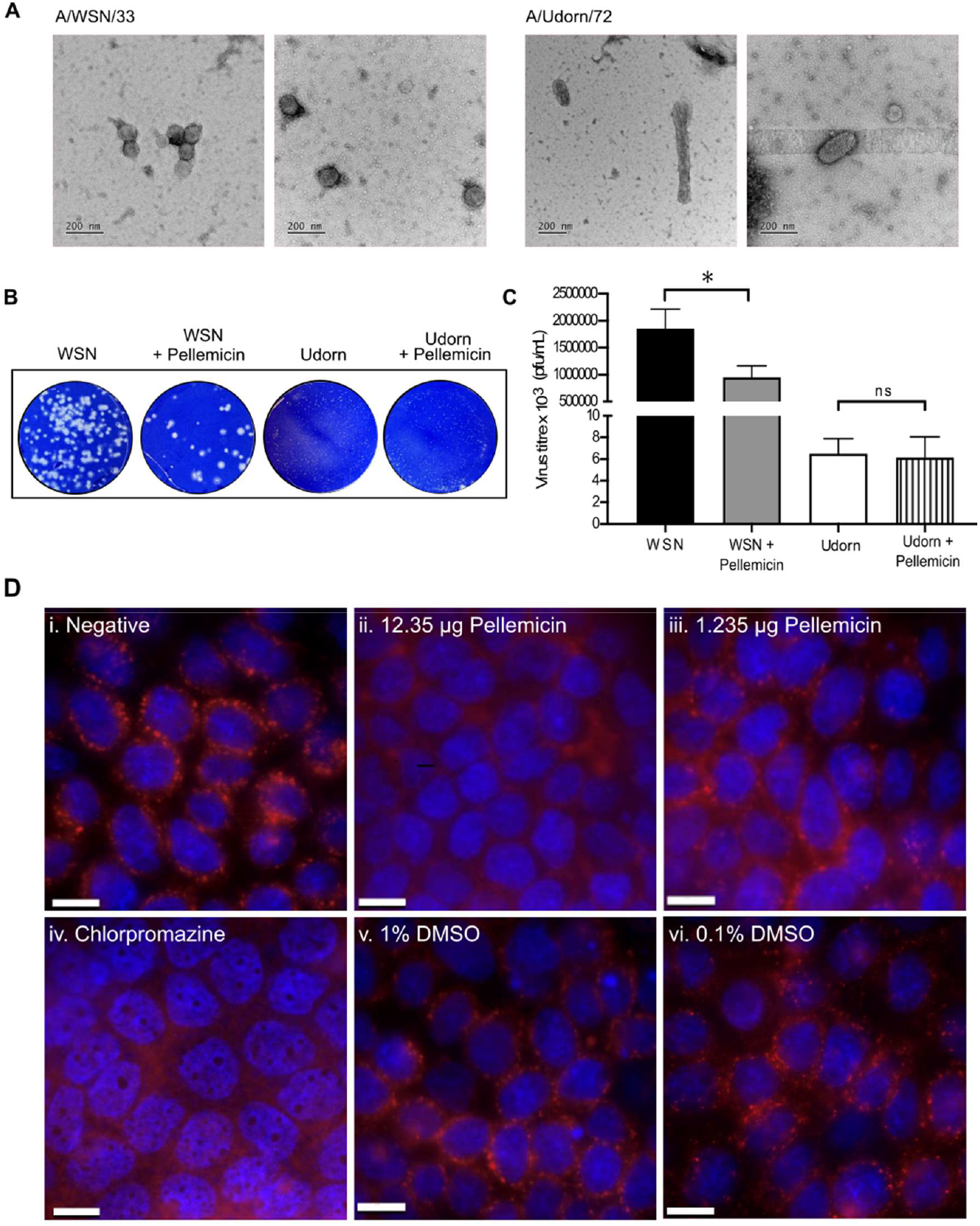
Pellemicin inhibits clathrin-mediated endocytosis. A) Electron microscope images of A/WSN/33 or A/Udorn/72 virus stocks. Scale bar 200 nm. B) MDCK cells were infected with influenza A virus A/WSN/33 (H1N1) (MOI 0.1) or A/Udorn/72 (H3N2) (MOI 0.01) and treated with pellemicin (0.1235µg/mL) or left untreated for 1 h. Representative images of plaque assays are shown. C) Mean viral titres of WSN alone (black), WSN with pellemicin (grey), Udorn alone (white) and Udorn with pellemicin (striped), from three experiments. Untreated viruses were compared to pellemicin-treated group using an unpaired t-test. * p<0.05 was considered statistically significant. Error bars show standard deviation. D) MDCK cells were grown on coverslips before being i) left untreated, ii) treated with 12.35 μg pellemicin, iii) treated with 1.235 μg pellemicin, iv) treated with 20 µM endocytosis inhibitor chlorpromazine (CPZ), v) treated with 1% DMSO, or vi) treated with 0.1% DMSO. Alexa-647N-conjugated transferrin was added to the cells for 1 hour before they were visualised with a widefield microscope. Scale bar 10 µm.

This result was further explored using transferrin as a specific marker for clathrin-mediated endocytosis. MDCK cells were grown on coverslips before being treated with 12.35 μg or 1.235 μg pellemicin or left untreated. Alexa-647N-conjugated transferrin was then added to the cells for 1 hour before being visualised with a widefield microscope. The untreated cells showed transferrin accumulation in distinct cytoplasmic puncta inside the cells, suggesting that transferrin was efficiently internalised (Fig 7D panel i). In contrast, treatment of the MDCK cells with 12.35 µg pellemicin significantly reduced transferrin internalisation (Fig 7D panel ii), whilst treatment with a lower concentration of 1.235 µg pellemicin also showed reduced cytoplasmic puncta compared to untreated cells (Fig 7D panel iii). A control where 20 μM chlorpromazine (CPZ), a clathrin-mediated endocytosis inhibitor, was used, showed a similar reduction in transferrin internalisation to pellemicin addition (Fig 7D panel iv), and DMSO only controls showed transferrin accumulation in cytoplasmic puncta as a result of endocytosis (Fig 7D panel v & vi). In summary, transferrin uptake into cells was decreased when the cells were pre-treated with pellemicin compared with the controls, indicating that pellemicin is an inhibitor of clathrin-mediated endocytosis and thus suggesting a mechanism of action of the compound against the influenza virus.

## Discussion

Spirotetronates were originally identified as antibiotics with potent activity against Gram-positive bacteria. However, several spirotetronates have since been shown to possess promising antiviral properties, although their mechanism(s) of antiviral action are poorly understood. Here, we report the identification of a putative spirotetronate BGC in the mycetoma pathogen *A. pelletieri* DSM 43383. Predictive sequence analyses of the enzymes encoded by the BGC led us to propose that it directs the assembly of a metabolite with an identical planar structure to MM46115 (here renamed pellemicin), a class II spirotetronate first reported in 1990 with promising activity against the parainfluenza virus.^15^ Despite extensive spectroscopic analyses,^16^ the relative stereochemistry of pellemicin could not be fully resolved and its absolute stereochemistry remained unknown. We showed that *A. pelletieri* DSM 43383 produces a metabolite with the same molecular formula and identical ^1^H and ^13^C NMR spectra as pellemicin. Exploiting a combination of established sequenced-based tools for predicting the stereochemical outcome of reactions catalysed by modular PKS KR, DH, and ER domains,^21, 22, 24^ our recent discovery and stereochemical characterisation of diene-forming DH domains in modular PKSs,^23^ and innovative phylogenetic approaches to predicting the stereocontrol imparted by spirotetronate-forming Diels-Alderases, we confirmed the previously reported relative stereochemical assignments for pellemicin,^16^ in addition to resolving remaining relative and absolute stereochemical ambiguities. Comparative analysis of the NMR spectroscopic data for pellemicin and *S*-*exo, S*-*endo* and *R*-*endo* type II spirotetronates that have been structurally elucidated by X-ray crystallography, combined with comparison of the measured ECD spectrum for pellemicin with that calculated for its aglycone, provided unambiguous experimental support for our stereochemical assignments.

Our data reveal pellemicin is an extremely rare member of the spirotetronate family that appears to arise from intramolecular [4+2] cycloaddition of a conjugated *E, Z*-diene to the *re*-face of the 2-acyl-4-methylenetetronate via an *endo* transition state. The recently reported wychimicins are the only other type II spirotetronates that appear to result from a cycloaddition with these stereochemical features, giving rise to a common 20*S*, 23*S*, 2′*R* configuration with pellemicin.^27^ The wychimicins were proposed to be the first members of a new family of class II spirotetronates.^27^ However, our work shows that pellemicin, first reported over three decades ago,^15^ is in fact the founding member of this unusual family. We recently identified the putative wychimicin biosynthetic gene cluster in the draft genome sequence of *Actinocrispum wychmicini* DSM 45934 and preliminary sequence analysis indicates many similarities between the pathways for assembly of pellemicin and the wychimicins, alongside some unique elements that explain the structural differences (L.A.M.M. Natalie To, and G.L.C., unpublished data). The discovery of type II spirotetronates arising from [4+2] cycloadditions following distinct stereochemical trajectories to well-characterised examples resulting in S-*exo* and S-*endo* adducts (e.g. chlorothricin and pyrrolosporin, respectively) provides an opportunity to better understand structure-activity relationships within this fascinating group of polyketides.

We also demonstrated that pellemicin inhibits both influenza A and influenza B viruses at concentrations which are not toxic to mammalian cells. Plaque assays showed that pellemicin demonstrated anti-influenza activity with an IC_50_ of 0.34 µg/mL. We then examined the cytotoxic properties of pellemicin on healthy cells using both a neutral red cell cytotoxicity assay to measure cell viability and a colorimetric cell proliferation assay based on the incorporation of thymidine during DNA synthesis in replicating cells. Data from both assays showed that addition of pellemicin at of 0.12 µg/mL did not significantly reduce cell viability, indicating that pellemicin can produce antiviral activity against the influenza virus without cytotoxicity. This differs from the results of the Ashton et al.^15^ paper, which used a similar thymidine incorporation assay and found that 0.41 µg pellemicin was toxic to MDCK cells, in comparison to our results which suggest that up to 12.35 µg showed no significant cytotoxicity. As our results obtained by the cell viability and proliferation assays were in agreement, the discrepancy with the previous study may be due to their visual interpretation of results, or the significance thresholds used to define cytotoxicity.

Pellemicin treatment of MDCK cells was shown to inhibit influenza A-induced cell death for up to 6 hours before virus titres reached untreated levels. We speculated that this was because 0.12 µg/mL of pellemicin did not completely inhibit viral growth and a low level of viral replication still occurs. As a single cycle of viral replication takes approximately 6 hours, new virus particles will be released from infected cells after this timepoint, resulting in an increase in viral titre.^33^ We confirmed this by adding pellemicin into the experiment again at 6 hours post-infection, and showed that viral titres remained low. Although the precise stability of pellemicin is unknown, this suggests that the compound is metabolised by the cell within 6-8 hours, as there is no stable compound remaining to inhibit newly made virus particles.

Pre- and post-incubation assays showed that pellemicin has antiviral activity when added either together or before virus addition but had no effect if added post-infection, once the virus has already entered the cells. Inclusion of a wash step after pellemicin addition but before infection did not reduce the antiviral activity of the compound, suggesting that pellemicin is internalised into the cell. These results, combined with haemagglutinin, neuraminidase and qRT-PCR assays, suggested that pellemicin is inhibiting an early step of the viral life cycle but that it does not block binding of viral HA to host receptors to cause antiviral activity. Further evidence for the mechanism of action of pellemicin was provided by the observation that addition of the compound inhibited clathrin-mediated endocytosis of transferrin in a concentration dependent manner, thereby providing a possible explanation for how pellemicin inhibits influenza virus proliferation.

The influenza virus exploits multiple endocytic pathways for infection, with clathrin-mediated endocytosis likely to be the most common.^41^ Interestingly, another natural product, ikarugamycin, which was first isolated from cultures of *Streptomyces phaeochromogenes* and shown to have anti-protozoal activity, has also been shown to inhibit clathrin-mediated endocytosis in plant cells.^36^ Ikarugamycin contains a tetramate moiety, which has similarities to the spirotetronate in pellemicin. Tetronic acids are known to form chelates with various metal ions, in particular, tetronate containing natural products kijanimicin, quatromicin and tetrocarcin have been reported to strongly chelate metals.^37^ This leads us to speculate that pellemicin may be inhibiting endocytosis of viral particles by chelating calcium, given the importance of calcium influx in clathrin-mediated endocytosis of influenza A and many other viruses.^38^ This would be consistent with a general mechanism of action for spirotetronates that have been shown to display antiviral activity. The Ashton et al.^15^ study suggested that pellemicin displayed antiviral activity against parainfluenza, influenza and RSV (albeit with high cytotoxicity for the latter two), and hence whether pellemicin might act as a general inhibitor against other enveloped virus infections, especially by a functionally similar mode of action, deserves further investigation.

In conclusion, we identified the pellemicin BGC in the genome sequence of the mycetoma pathogen *A. pelletieri* DSM 43383 and demonstrated it is a novel producer of this unusual spirotetronate. Using a combination of known and novel bioinformatics-based approaches, we were able to predict and experimentally verify the relative and absolute stereochemistry of pellemicin. While the value of modular PKS KR, DH, and ER domain predictive analysis has been demonstrated as a tool for accurate stereochemical prediction of linear polyketide chains,^38,39,40^ the work described herein highlights the utility of our recent discovery and stereochemical characterisation of diene-forming DH domains to extend such approaches.^23^ Moreover, the phylogeny-based prediction of stereochemical outcome for spirotetronate-forming Diels-Alder reactions reported here showcases the potential of predictive bioinformatics to be employed in tackling more complex stereochemical challenges, such as elucidating the relative and absolute stereochemistry of polycyclic ring systems. Pellemicin was shown to effectively inhibit the proliferation of influenza viruses without significant cytotoxicity. Subsequent mechanistic studies indicated that pellemicin acts at an early stage of the viral life cycle and is likely to inhibit influenza virus entry into the cell by inhibiting endocytosis. It is possible that pellemicin could display a broad-spectrum antiviral activity that may be effective against a variety of enveloped viruses. Further investigations of pellemicin biosynthesis will enable the development of bioengineering approaches to the production of structural analogues that could provide valuable insights into the structure-activity relationship and may ultimately lead to a new class of antivirals with therapeutic potential.

## Resource availability

### Lead contact

Further information and resources should be directed to and will be fulfilled by the lead contact, Nicole Robb.

### Materials availability

Materials generated in this study will be made available on request, but we require a completed materials transfer agreement if there is potential for commercial application. All requests for resources and reagents should be directed to the lead contact.

## Data and code availability

All data reported in this paper will be shared by the lead contact upon request.

## Acknowledgments

The authors wish to thank Dr. Saskia Bakker for assistance with the negative stain electron microscopy, as well as Drs. Andrew Bottrill and Cleidi Zampronio from The University of Warwick’s Proteomics Facility, and Dr. Maëlle Lorvellec of CAMDU (Computing and Advanced Microscopy Unit) for their support & assistance in this work. Dr Ivan Prokes and Dr Lijiang Song are thanked for assistance with acquisition of NMR and LC-MS data, respectively. This work was supported by a Royal Society Dorothy Hodgkin Research Fellowship DKR00620 and Research Grant for Research Fellows RGF\R1\180054 (to N.C.R.), the MRC-Interdisciplinary Biomedical Research Doctoral Training Partnership (grant ref. MR/N014294/1; studentship to A.T.), the BBSRC Midlands Integrative Biosciences Doctoral Training Partnership (grant refs. BB/M01116X/1, NE/W503095/1, and BB/T00746X/1; studentships to R.A.C., M.C.H.B.D. and C.F.U.), and a BBSRC responsive mode grant and China Partnering Award (BB/S008020/1 and BB/L010852/1, respectively, to G.L.C.). L.A.M.M. was supported by an Australian Research Council Discovery Early Career Researcher Award (DE240100502). Y.H. and N.L. were funded by the Strategic Priority Research Program of the Chinese Academy of Sciences (XDB0810000), the National Natural Science Foundation of China (No. 81773615 and 31670502), the International Partnership Program of Chinese Academy of Sciences (153211KYSB20170014).

## Author contributions

Conceptualisation, L.M.A.; G.L.C.; N.C.R.; Methodology, A.T.; R.A.C.; L.M.A.; G.L.C.; N.C.R.;

Investigation, A.T.; L.A.M.M.; R.A.C.; C.F.U.; J.Z.; S.C.L.; M.C.H.B.D.; N.L.; Writing – original draft, A.T.; L.A.M.M.; R.C. Writing – review & editing, N.C.R.; G.L.C; Funding acquisition, L.A.M.M.; L.M.A.; G.L.C.; N.C.R.; Resources, C.P.T.; Supervision, Y.H.; L.M.A.; G.L.C.; N.C.R.

## STAR Methods

### Key resources table

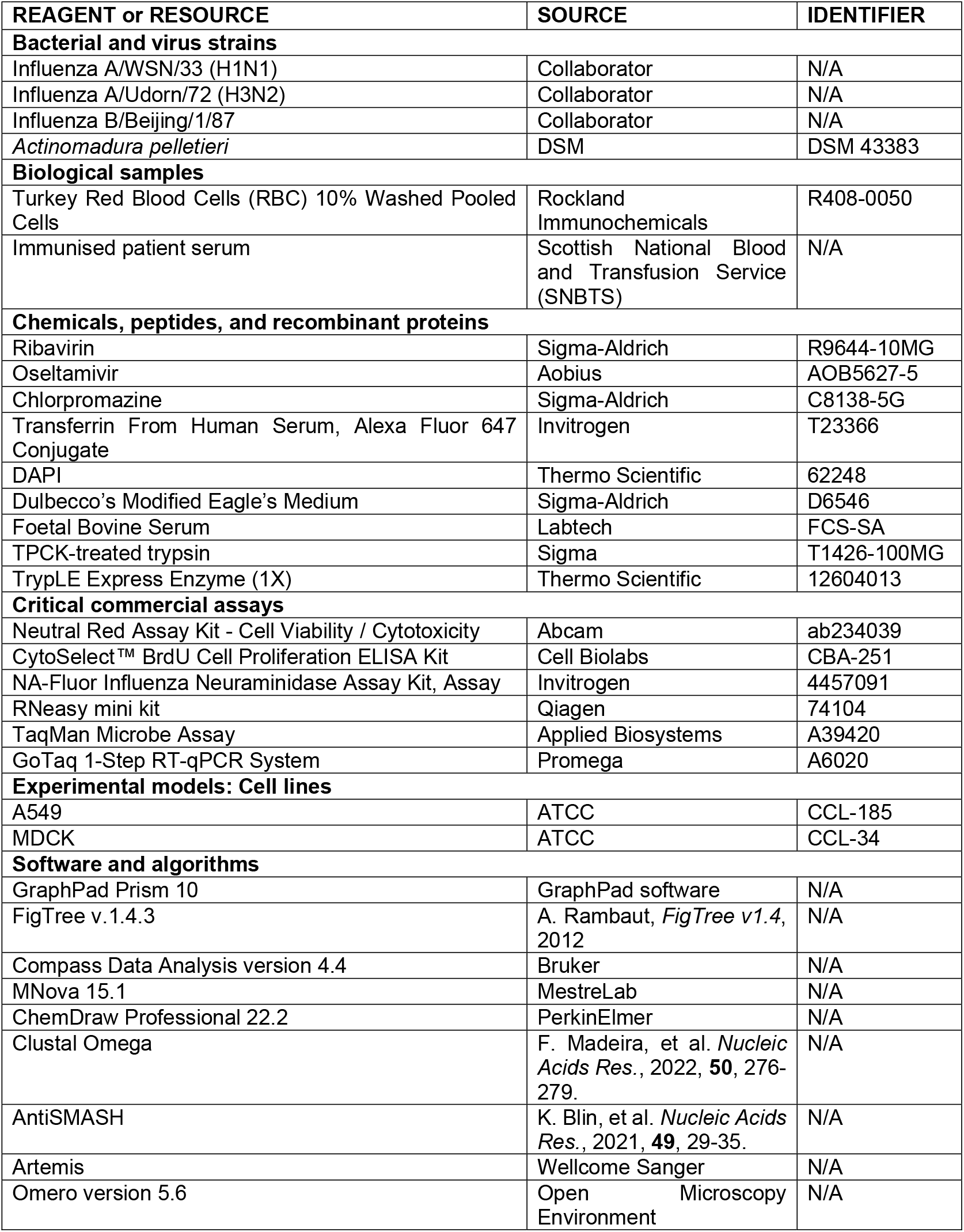

| REAGENT or RESOURCE | SOURCE | IDENTIFIER |
| --- | --- | --- |
| **Bacterial and virus strains** |  |  |
| Influenza A/WSN/33 (H1N1) | Collaborator | N/A |
| Influenza A/Udorn/72 (H3N2) | Collaborator | N/A |
| Influenza B/Beijing/1/87 | Collaborator | N/A |
| *Actinomadura pelletieri* | DSM | DSM 43383 |
| **Biological samples** |  |  |
| Turkey Red Blood Cells (RBC) 10% Washed Pooled Cells | Rockland Immunochemicals | R408-0050 |
| Immunised patient serum | Scottish National Blood and Transfusion Service (SNBTS) | N/A |
| **Chemicals, peptides, and recombinant proteins** |  |  |
| Ribavirin | Sigma-Aldrich | R9644-10MG |
| Oseltamivir | Aobius | AOB5627-5 |
| Chlorpromazine | Sigma-Aldrich | C8138-5G |
| Transferrin From Human Serum, Alexa Fluor 647 Conjugate | Invitrogen | T23366 |
| DAPI | Thermo Scientific | 62248 |
| Dulbecco's Modified Eagle's Medium | Sigma-Aldrich | D6546 |
| Foetal Bovine Serum | Labtech | FCS-SA |
| TPCK-treated trypsin | Sigma | T1426-100MG |
| TrypLE Express Enzyme (1X) | Thermo Scientific | 12604013 |
| **Critical commercial assays** |  |  |
| Neutral Red Assay Kit - Cell Viability / Cytotoxicity | Abcam | ab234039 |
| CytoSelect™ BrdU Cell Proliferation ELISA Kit | Cell Biolabs | CBA-251 |
| NA-Fluor Influenza Neuraminidase Assay Kit, Assay | Invitrogen | 4457091 |
| RNeasy mini kit | Qiagen | 74104 |
| TaqMan Microbe Assay | Applied Biosystems | A39420 |
| GoTaq 1-Step RT-qPCR System | Promega | A6020 |
| **Experimental models: Cell lines** |  |  |
| A549 | ATCC | CCL-185 |
| MDCK | ATCC | CCL-34 |
| **Software and algorithms** |  |  |
| GraphPad Prism 10 | GraphPad software | N/A |
| FigTree v.1.4.3 | A. Rambaut, *FigTree v1.4*, 2012 | N/A |
| Compass Data Analysis version 4.4 | Bruker | N/A |
| MNova 15.1 | MestreLab | N/A |
| ChemDraw Professional 22.2 | PerkinElmer | N/A |
| Clustal Omega | F. Madeira, et al. *Nucleic Acids Res.*, 2022, **50**, 276-279. | N/A |
| AntiSMASH | K. Blin, et al. *Nucleic Acids Res.*, 2021, **49**, 29-35. | N/A |
| Artemis | Wellcome Sanger | N/A |
| Omero version 5.6 | Open Microscopy Environment | N/A |

## Experimental models

### Cells and viruses

Madin-Darby canine kidney (MDCK) or A549 alveolar basal epithelial cells were grown to confluency in Dulbecco’s minimal essential medium supplemented with 10% foetal calf serum (DMEM/10% FBS), at 37°C, 5% CO_2_. For antiviral assays, confluent MDCK cells were trypsinised and 2mL cell solution seeded in 6-well plates before incubating at 37°C, 5% CO_2_ overnight. At confluency, these plates contain ∼1 x 10^6^ cells per well. For cytotoxicity assays, 24-well plates with 1 mL cell solution (∼0.25 x 10^6^ cells) per well or 96-well plates with 0.5 mL cell solution (∼0.05 x 10^6^ cells) per well were used.

Influenza A strains H1N1 (A/WSN/33) and H3N2 (A/Udorn/72), and influenza B (B/Beijing/1/87) strains were used. To culture the viruses, confluent flasks of MDCK cells were infected with the virus stock (1 mL) and, for A/Udorn/72 and B/Beijing/1/87, supplemented with 2 µg/mL TPCK-treated trypsin. Flasks were incubated for 1 h at 37°C, 5% CO_2_. 9 mL DMEM/0.5% FBS was added, and the solution was incubated for 3 days at 37°C, 5% CO2. The culture medium was clarified by centrifugation at 1000 rpm for 5 min. The supernatant containing the virus was collected, aliquoted and stored at -80°C. Virus stocks were titred by plaque assay in MDCK cells. Virus infections were carried out at a multiplicity of infection (MOI) of 0.1.

## Method details

### Bioinformatic methods

AntiSMASH was used to identify gene clusters from genomic DNA, as well as to identify individual genes within the sequence of a gene cluster.^42^ Gene clusters were visualised in Artemis.^43^ Functions of the enzymes encoded by the genes were identified by sequence comparison to homologous enzymes in the NCBI BLAST database.^44^ Sequence alignments were generated using Clustal Omega.^45^ For phylogenetic analysis, sequences of spirotetronate forming Diels-Alderases were identified using BLASTp. A multiple sequence alignment was performed using ClustalX 2.1. A phylogenetic tree was generated using the Neighbour-Joining algorithm on the Diels-Alderase multiple sequence alignment in FigTree v.1.4.390 software.

### Pellemicin production

Experiments were performed using a glycerol stock of *A. pelletieri* (DSM 43383). Stocks were produced by culturing the bacterial strain in 50 mL liquid International Streptomyces Project 2 (ISP2) medium (0.4% yeast extract powder, 1% malt extract powder, 0.4% glucose (with 2% agar in solid medium) in distilled water, pH 7.2 before sterilisation) for 7 days (with shaking at 240 rpm, 28°C) and adding 50% glycerol (1:1 volume) to the culture. *A. pelletieri* glycerol stock (0.04%) was spread on a plate containing GYM Streptomyces medium (0.4% yeast extract, 1% malt extract, 0.2% calcium carbonate, 0.4% glucose (with 2% agar in solid medium) in distilled water, pH 7.2 before sterilisation). The plate was incubated for 10 days at 28°C. Red colonies were transferred from the agar plate to 50 new agar plates and incubated for 10 days, at 28 °C. To extract pellemicin, ethyl acetate was added to the bacterial culture (1:1 volume), followed by shaking every 10 min for 1 h. The solvent layer was then removed using a rotary evaporator at 200 rpm and 30°C (Rotavapor R-300, Buchi) connected to a vacuum pump (V-700, Buchi) to obtain the dry extract.

### Reversed-phase high-performance liquid chromatography (HPLC)

5 mL methanol was added to the extract then centrifuged at 13,200 rpm for 10 min. Water and acetonitrile (1:1 ratio) were used as the mobile phase on the Agilent 1200 series, with a flow rate of 9 mL/min. The absorbance was measured at 210 nm and fractions collected every 2 min.

### Liquid chromatography-high resolution mass spectrometry (LC-HRMS)

Samples were analysed by ultrahigh-performance liquid chromatography (UHPLC) using a Dionex 3000 UHPLC system connected to a C18 column (100 × 2.1 mm, 1.8 μm), coupled to a Bruker Compact ESI-Q-TOF mass spectrometer. The mass spectrometer was used in positive ion mode, using sodium formate (10 mM) for calibration. MS data was analysed using Compass Data Analysis software (version 4.4).

### NMR spectroscopic analysis of pellemicin

NMR spectroscopic analysis of purified pellemicin was carried out on a Bruker 500 MHz spectrometer equipped with a DCH cryoprobe in CDCl_3_ at 25 °C. The ^1^H and ^13^C NMR signals were referenced to the unlabelled solvent signal at 7.26 ppm (^1^H) and 77.16 ppm (^13^C). ^1^H, ^13^C (JMOD), COSY, HSQC, HMBC, NOESY, ROESY and TOCSY NMR experiments were used to confirm the structure of pellemicin. Spectra were processed with MestReNova software.

### Plaque assays

The virus stock was ten-fold serially diluted in DMEM/0.5% FBS. 500 μL of each serial dilution were added to each well of a confluent 6-well plate and kept on a shaker at room temperature for 1 h. The virus solution was aspirated before 2% agarose was diluted 1:1 with DMEM/0.5% FBS and added to each well. Plates were kept at room temperature to allow the overlay to solidify, before incubating at 37°C, 5% CO_2_, for 3 days. Coomassie Blue stain (2% Coomassie Blue powder, 7.5% acetic acid, 50% ethanol absolute, diluted in Milli-Q water) was used to stain live cells. Plaques were manually counted to calculate viral titres.

### Neutral red cell viability assay

Pellemicin (123.5 – 0.01235 μg), ribavirin (5 – 0.625 μg) and DMSO (1 – 0.0001%) were prepared in DMEM/0.5% FBS. 250 μL test dilutions were added in triplicate to confluent 24-well plates and incubated for 2 h (37°C, 5% CO_2_). DMEM/0.5% FBS (750 μL) was then added to wells and incubated for 3 days. The Neutral Red Assay kit was used (ab234039, Abcam) to measure cell viability. After incubation, cells were washed with 1X washing solution and stained with 1X Neutral Red staining solution. The plate was incubated for 2 h, before removing the stain and adding 1X solubilisation solution. The plate was rocked at room temperature for 20 min, before using the SPECTROstar plate reader (BMG Labtech) to measure absorbance at 540 nm. Mean absorbance was calculated and the DMSO background removed from the pellemicin and ribavirin conditions. Absorbance of cells in DMEM/0.5% FBS was set at 100% viability and test conditions were normalised to this value.

### Cell proliferation assay

Pellemicin (123.5 – 0.01235 μg) and DMSO (1 – 0.0001%) were prepared in DMEM/0.5% FBS. 100 μL test dilutions were added in triplicate to confluent 96-well plates and incubated for 2 h (37°C, 5% CO2). DMEM/0.5% FBS (400 μL) was then added to cells and incubated for 3 days. The CytoSelect BrdU cell proliferation ELISA kit (Cell Biolabs Inc.) was used. 10X BrdU solution was added to wells and incubated for 3 h (37°C, 5% CO2). Cells were washed with PBS and fixed with Fix/Denature solution for 30 min. Anti-BrdU antibody was added for 1 h at room temperature, before washing with 1X Wash Buffer and replacing with Secondary Antibody HRP conjugate. Stop solution was used to end the reaction and absorbance was measured at 450 nm with Agilent BioTek Cytation 5 reader. Mean absorbance was calculated and the DMSO background removed from the pellemicin conditions. Absorbance of cells in DMEM/0.5% FBS was used as the negative control.

### Haemagglutination assay

Turkey red blood cells (RBCs) were supplied as a 10% suspension in PBS (Rockland Immunochemicals). Ten-fold serial dilutions of influenza A/WSN/33 stock in DMEM/0.5% FBS were added to 96-well V-bottom plates before a 1% RBC solution (diluted in PBS) was added to each well and incubated for 30 min. When pellemicin was added, dilutions of the compound were prepared in PBS and mixed with the virus before adding to the wells. Two-fold dilutions of serum from a patient immunised with the influenza vaccine were included as a positive control, with and without virus. Control groups with pellemicin (123.5 – 0.1235 µg), DMSO (1-0.001%) and PBS alone (negative) were also included.

### Neuraminidase inhibition assay

The NA-Fluor Influenza Neuraminidase Assay Kit (Thermo Fisher) was used. Pellemicin (100-0.0001 µM) or oseltamivir (10-0.00001 µM) were prepared in PBS before being added to A/WSN/33 virus or 1X assay buffer without virus, in a black 96-well plate. The plate was incubated for 30 min at 37°C before adding 200 µM NA-Fluor 2′-(4-methylumbelliferyl)-α-d-*N*-acetylneuraminic acid (MUNANA) substrate to each well and incubating for 1 h on a shaker at 37°C. 100 µL NA-Fluor stop solution was added and the plate was read on an Agilent BioTek Cytation 5 reader (excitation 350 nm and emission 440 nm). Background fluorescence of pellemicin and oseltamivir without virus was subtracted from values with virus.

### Real-time quantitative reverse transcription-PCR

Confluent 6-well plates of MDCK cells were infected with A/WSN/33 alone or with pellemicin (0.1235 µg) for 2-24 h. The cells were then trypsinised with 1X TryplE Express Enzyme, before DMEM/0.5% FBS was added to inactivate the trypsin. Samples were centrifuged (300 x g, 5 min) and supernatants and cell lysates collected separately. RNA extraction was performed with RNeasy mini kit (Qiagen). PCR was performed using the GoTaq Probe 1 Step RT-qPCR system (Promega). Target RNA sequences were identified using the primers and probes in the TaqMan Microbe Assay (A39420, Applied Biosystems). The kit included a plasmid containing target sequences for influenza A viral RNA which was used as a positive control. Nuclease-free water was used as a negative control. Cycling conditions for real-time PCR were as follows: 25°C for 2 min, reverse transcription at 50°C for 10 min, and denaturation of RT polymerase at 95°C for 2 min, followed by 40 cycles of 95°C for 15 s and 60°C for 1 min. PCR was conducted using the CFX Opus 384 detection system, and the data were analyzed using the BR.io application (Bio-Rad). Cq values were normalised to the negative control to calculate mean fold-change.

### Electron microscopy

Samples were fixed with 4% paraformaldehyde, adsorbed onto carbon coated formvar grids, negatively stained with 1% aqueous uranyl acetate and imaged in a JEOL 2100 Plus microscope operating at 200 kV.

### Fluorescence imaging

MDCK cells were grown on 13 mm coverslips in 24-well plates before being treated with 12.35 μg or 1.235 μg pellemicin for 2 hours at 37 °C. 20 μM chlorpromazine, a clathrin-mediated endocytosis inhibitor, was used as a positive control. DMSO vehicle controls for pellemicin and a 0.5% FBS/DMEM negative control were also included. The cells were then labelled with Alexa Fluor 647N-conjugated transferrin (25 μg/mL) for 1 hour at 4 °C to allow transferrin uptake. 4% paraformaldehyde was used to fix the cells for 15 min at room temperature, before the coverslips were mounted onto glass microscope slides with Mowiol containing DAPI (1 μL/mL). Cells were viewed using a widefield deconvolution microscope from Applied Precision Instruments (API) and images processed with Omero.

### Statistical analysis

All experiments were conducted in duplicate or triplicate, and mean values were calculated from a set of three individual repeats. All results are displayed as mean ± standard deviation. Statistical analyses were performed with GraphPad Prism 10 and statistical significance was calculated using the unpaired Student’s *t* test, or one-way ANOVA. IC50 values were calculated using linear regression analysis to generate a line of best fit. The statistical test used for each experiment is indicated in the respective figure legend. p < 0.05 was considered significant across all experiments.

