## Supplementary material for "Bioinformatics-based stereochemical elucidation of MM 46115: an unusual antiviral spirotetronate that inhibits clathrin-mediated endocytosis"

### I. Biosynthetic and Stereochemical Elucidation of pellemicin

#### Pellemicin genes in the biosynthetic gene cluster

**Table S1.** Proposed functions of proteins encoded by the pellemicin BGC in *A. pelletieri* DSM 43383

| Gene BZB76_ | Protein size (aa) | Proposed function | homologue in *chl* BGC | Origin of nearest homologue | Identity/ similarity (%) | Accession No. |
|---|---|---|---|---|---|---|
| **6430** | 429 | Transporter (CPA2 family) | | *Actinomadura montaniterrae* | 64/76 | WP_151541937.1 |
| **6431** | 461 | dNDP-4-dehydro-6-deoxy-α-D-glucose-2,3-dehydratase | ChlC3 | *Streptomyces lucensis* | 65/73 | WP_22981 6081.1 |
| **6432** | 79 | Hypothetical protein | | *Actinomadura geliboluensis* | 54/64 | WP_13863 4234.1 |
| **6433** | 262 | Type II Thioesterase | | *Actinomycetia bacterium* | 68/77 | MCL2551318.1 |
| **6434** | 266 | DNA-binding SARP family transcriptional activator | ChlF2 | *Streptomyces sulfonofaciens* | 72/81 | WP_18993 8827.1 |
| **6435** | 754 | RND superfamily putative drug exporter | | *Streptomyces* sp. *RTd22* | 56/72 | WP_063737295.1 |
| **6436** | 368 | Exomethylene tetronate synthase | ChlD4 | *Streptomyces* sp. ODS25 | 67/78 | WP_25595 5326.1 |
| **6437** | 266 | Acetyl transferase | ChlD3 | *Actinocrispum wychmicini* | 71/81 | WP_132116034.1 |
| **6438** | 754 | ACP | | *Actinomadura geliboluensis* | 57/77 | WP_138634256.1 |
| **6439** | 636 | *D*-Glyceryl-ACP synthase | ChlD1 | *Micromonospora* sp. *RP3T* | 65/76 | WP_107154962.1 |
| **6440** | 343 | Ketoacyl-ACP synthase | ChlM | *Actinocrispum wychmicini* | 76/87 | WP_13211 6040.1 |
| **6441** | 501 | Diels-Alderase (decalin) | ChlE3 | *Streptomyces armeniacus* | 61/71 | WP_20887 6620.1 |
| **6442** | 1507 | Type I modular PKS | | *Actinomadura geliboluensis* | 60/68 | TMR4177 2.1 |
| **6444** | 3910 | Type I modular PKS | | *Actinomadura kijaniata* | 62/72 | ACB464710.1 |
| **6446** | 1787 | Type I modular PKS | | *Streptomyces barringtoniae* | 56/68 | WP_22862 6829.1 |
| **6448** | 5163 | Type I modular PKS | | *Streptomyces kanamyceticus* | 52/64 | QEU900810.1 |
| **6450** | 1542 | Type I modular PKS | | *Streptomyces rhizosphaericus* | 54/65 | WP_164427656.1 |
| **6452** | 4403 | Type I modular PKS | | *Allokutzneria* sp. NRRL B-24872 | 49/61 | WP_08682 4394.1 |
| **6453** | 1835 | Type I modular PKS | | *Goodfellowiella coeruleoviolacea* | 57/67 | WP_25378 0133.1 |
| **6455** | 355 | α-*D*-Glucose-1-phosphate thymidylyl transferase | ChlC1 | *Streptomyces sulfonofaciens* | 65/77 | WP_189932193.1 |
| **6456** | 584 | 3-hydroxybutyryl-CoA dehydrogenase | | *Streptomyces* sp. W1SF4 | 60/68 | WP_12582 4799.1 |
| **6457** | 364 | β-Ketoacyl-ACP synthase | | *Streptomyces sabulosicollis* | 75/82 | WP_198269555.1 |
| **6458** | 444 | Crotonyl-CoA carboxylase/ reductase | | Kibdelosporangium persicum | 82/90 | NRN684040.1 |
| **6459** | 89 | ACP | ChlB2 | *Streptomyces antibioticus* | 51/67 | AAZ776750.1 |
| **6460** | 453 | $FADH_2$ and $O_2$-dependent halogenase | ChlB4 | *Actinocrispum wychmicini* | 79/88 | WP_132116064.1 |
| **6461** | 1785 | 6-methylsalicylic acid synthase | ChlB1 | *Streptomyces armeniacus* | 65/73 | WP_20887 8173.1 |
| **6462** | 347 | Ketoacyl-ACP synthase | ChlB3 | *Actinocrispum wychmicini* | 74/84 | WP_13211 6060.1 |
| **6463** | 67 | Hypothetical protein | | | | |
| **6464** | 186 | Diels-Alderase (spirotet) | ChlL | *Actinocrispum wychmicini* | 48/61 | WP_13211 6074.1 |
| **6465** | 384 | Aminotransferase | | *Streptomyces Exfoliates* | 69/81 | WP_030554129.1 |
| **6466** | 348 | Ketoacyl-ACP synthase | ChlB3 | *Amycolatopsis palatopharyngis* | 51/69 | WP_11605 1422.1 |
| **6467** | 406 | Glycosyltransferase | ChlC7 | *Actinocrispum wychmicini* | 48/65 | WP_13211 6056.1 |
| **6468** | 405 | Cytochrome P450 | | *Streptomyces reniochalinae* | 65/75 | WP_114017402.1 |
| **6469** | 425 | Signal transduction histidine kinase | | *Actinomadura sp. HBU206391* | 65/77 | WP_26198 7125.1 |
| **6470** | 505 | Hypothetical protein | | | | |
| **6471** | 242 | ABC-2 type transport system ATP-binding protein | | *Clostridium manihotivorum* | 43/64 | WP_238475790.1 |
| **6472** | 245 | dTDP-3-amino-3,4,6-trideoxy-α-*D*-glucopyranose *N*, *N*-dimethyltransferase | | *unclassified streptomyces* | 61/75 | WP_05110 4621.1 |
| **6473** | 372 | Aminotransferase | | *Streptomyces umbrinus* | 80/86 | WP_189842318.1 |
| **6474** | 327 | dTDP-α-*D*-glucose-4,6-dehydratase | | *Streptomyces lavendulae* | 72/83 | WP_03023 7981.1 |
| **6475** | 69 | Hypothetical protein | | | | |
| **6476** | 222 | LuxR family two-component transcriptional regulator | | *Dactylosporangium matsuzakiense* | 77/84 | WP_261962890.1 |
| **6477** | 972 | LuxR family transcriptional activator | | *Actinomadura geliboluensis* | 45/59 | WP_138641290.1 |

#### Sequence alignment of PKS domains

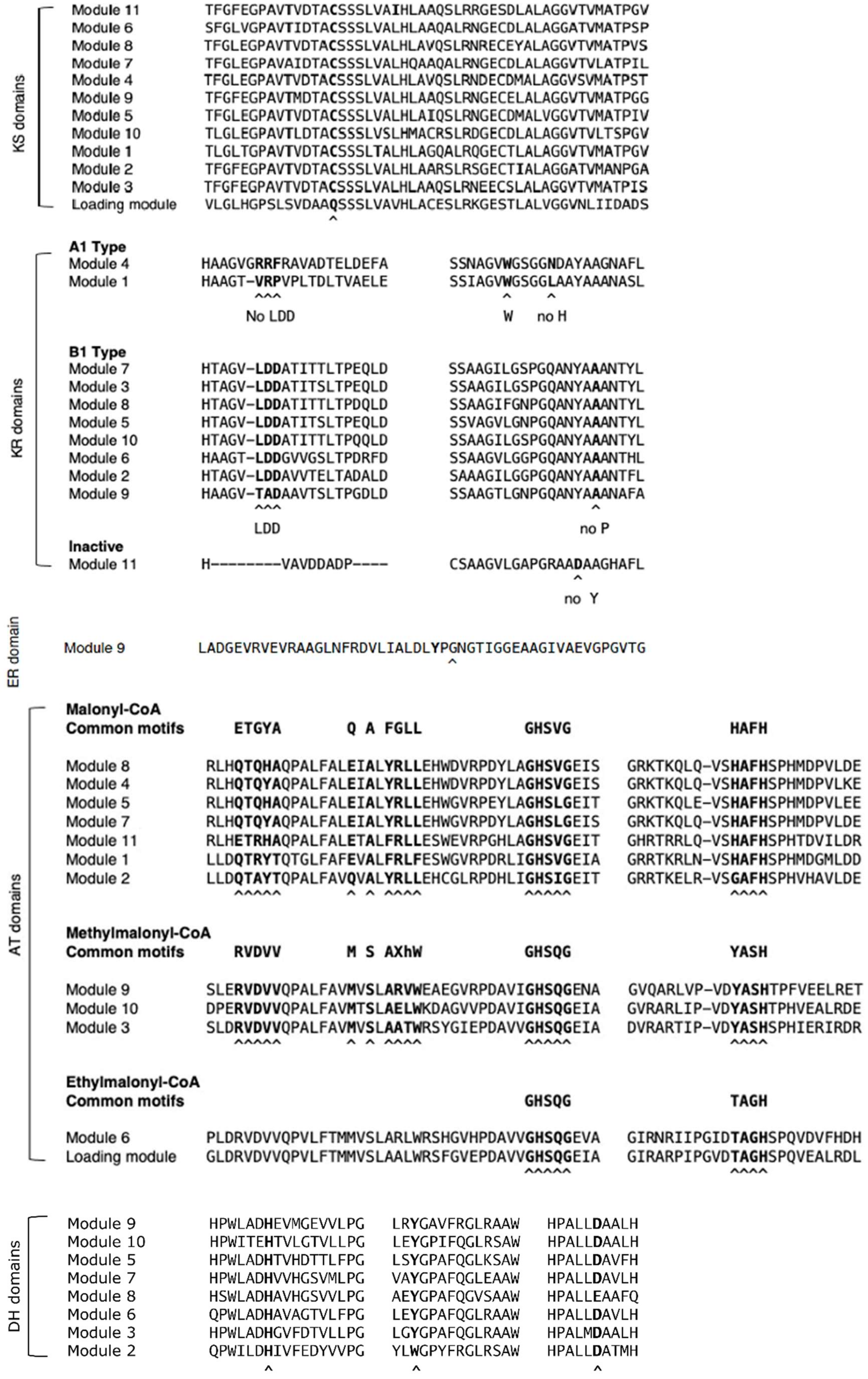

**Figure S1. Sequence alignment for each domain.** Relevant residues are highlighted in bold with ^ underneath.

#### Proposed mechanism for *E*, *Z*-diene formation

Full page width figure

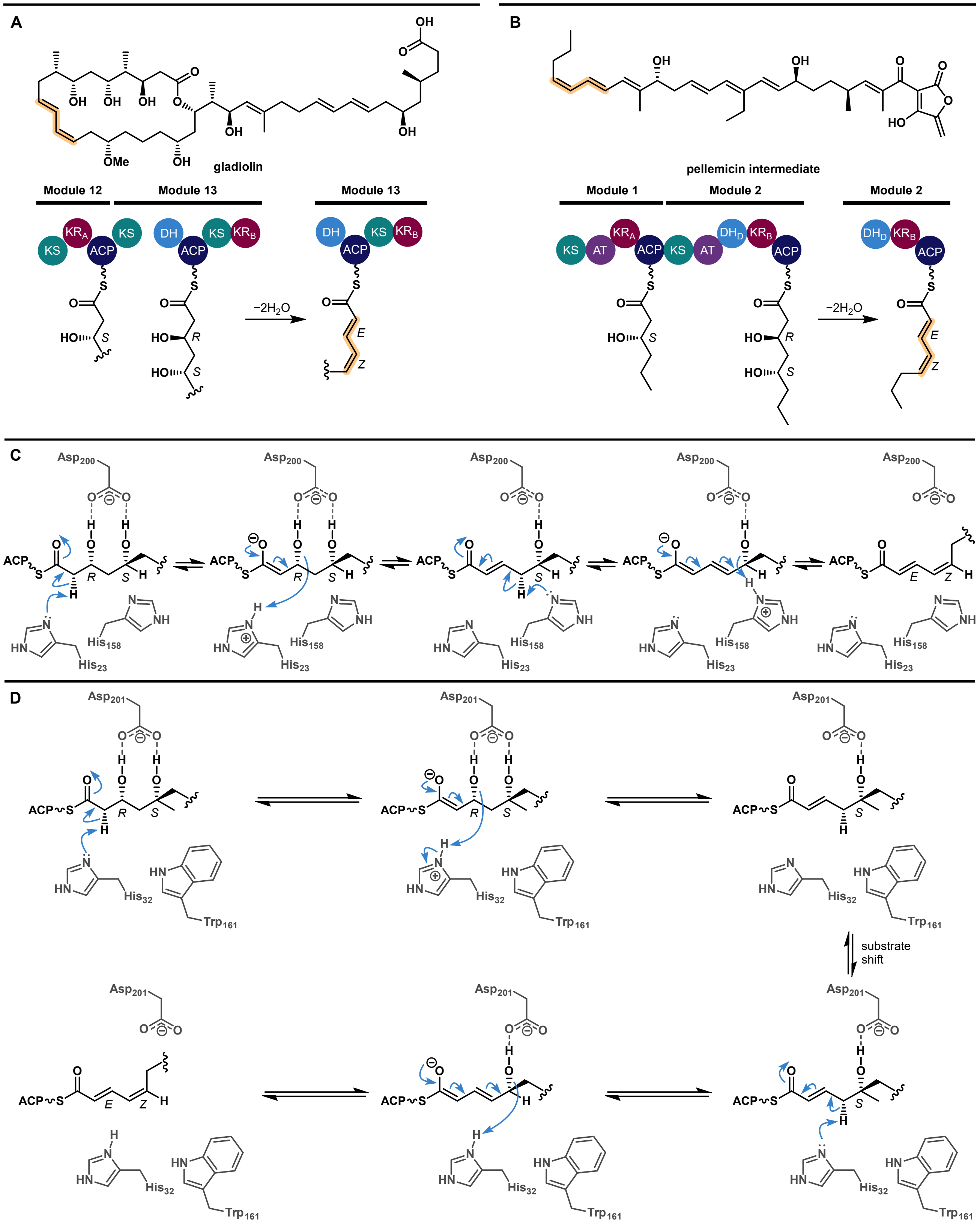

**Figure S2. Comparison of *E*, *Z*-diene formation in gladiolin and pellemicin biosynthesis.** A) Structure of gladiolin and *E*, *Z*-diene-forming step catalysed by the $DH^d$ domain in module 13 of the PKS. B) Structure of proposed product of the pellemicin PKS and *E*, *Z*-diene formation catalysed by the $DH^d$ domain in module 2 of the PKS. C) Proposed mechanism for *E*, *Z*-diene formation in gladiolin biosynthesis. D) Proposed mechanism for *E, Z*-diene formation in pellemicin biosynthesis.

#### Production and isolation of pellemicin

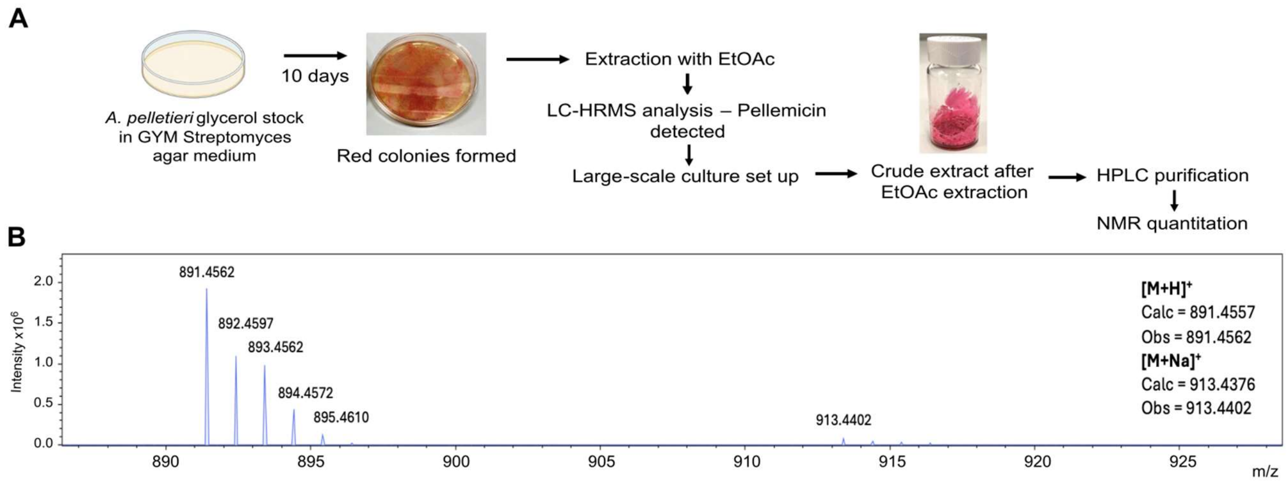

**Figure S3. Growth of *A. pelletieri* DSM 43383, UHPLC-ESI-Q-ToF-MS analysis of EtOAc extracts and purification from large scale cultures.** A) *A. pelletieri* DSM 43383 was streaked onto GYM agar plates and incubated for 10 days at 28 °C. The resulting red culture was extracted with ethyl acetate and UHPLC-Q-ToF-MS analsyes showed the extract contained a compound with molecular formula corresponding to pellemicin. Reverse-phase HPLC was used to purify pellemicin from ethyl actetate extracts of large-scale GYM agar cultures. B) Mass spectrum from UHPLC-ESI-Q-ToF-MS analysis of culture extracts. An ions with *m/z* = 891.4562 and 913.4402 corresponding to the $[M+H]^+$ and $[M+Na]^+$ ions of pellemicin were observed.

#### NMR spectra for pellemicin isolated from *A. pelletieri* DSM 43383 ($CDCl_3$)

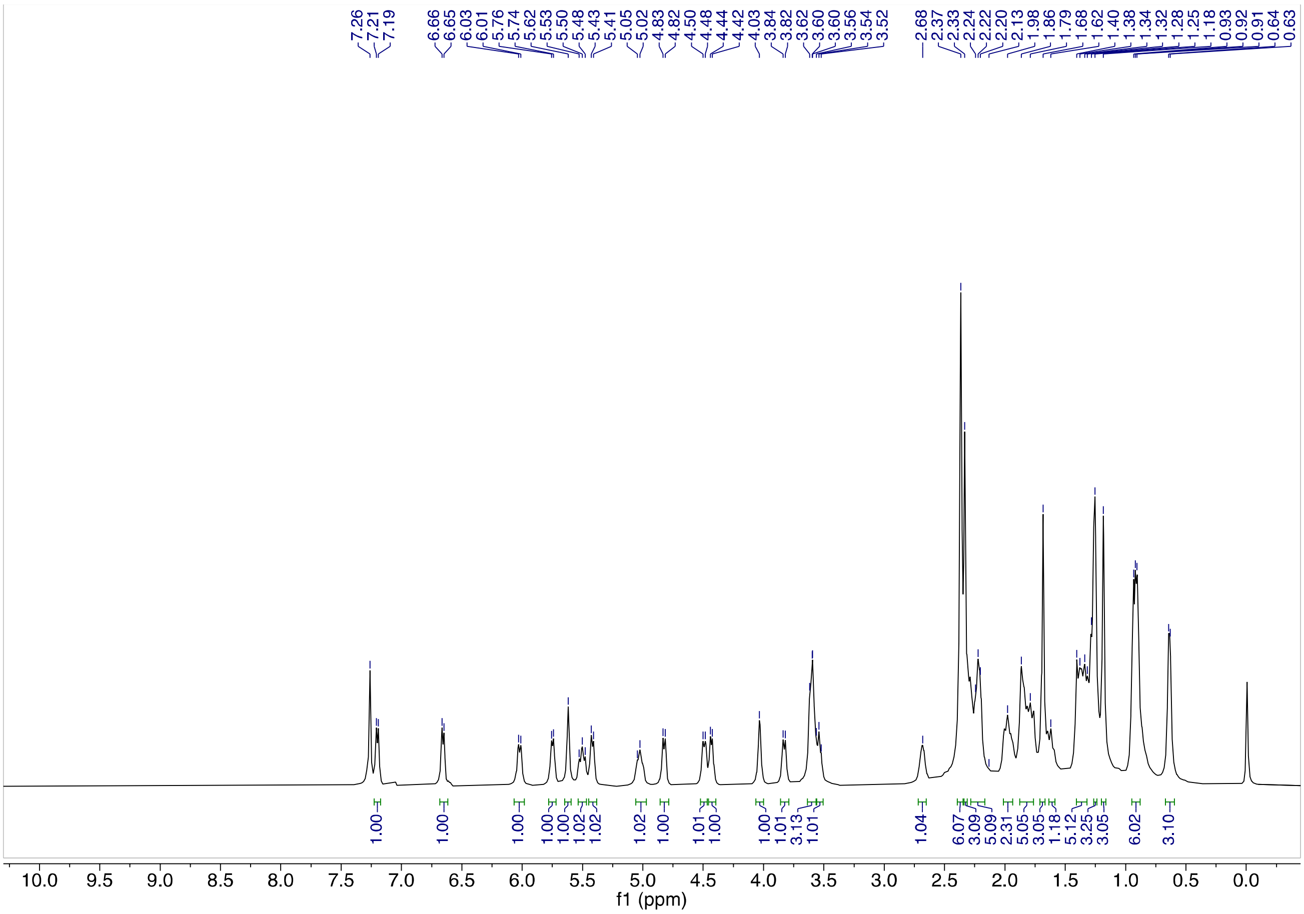

**Figure S4. ¹H NMR spectrum (500 MHz) of pellemicin.**

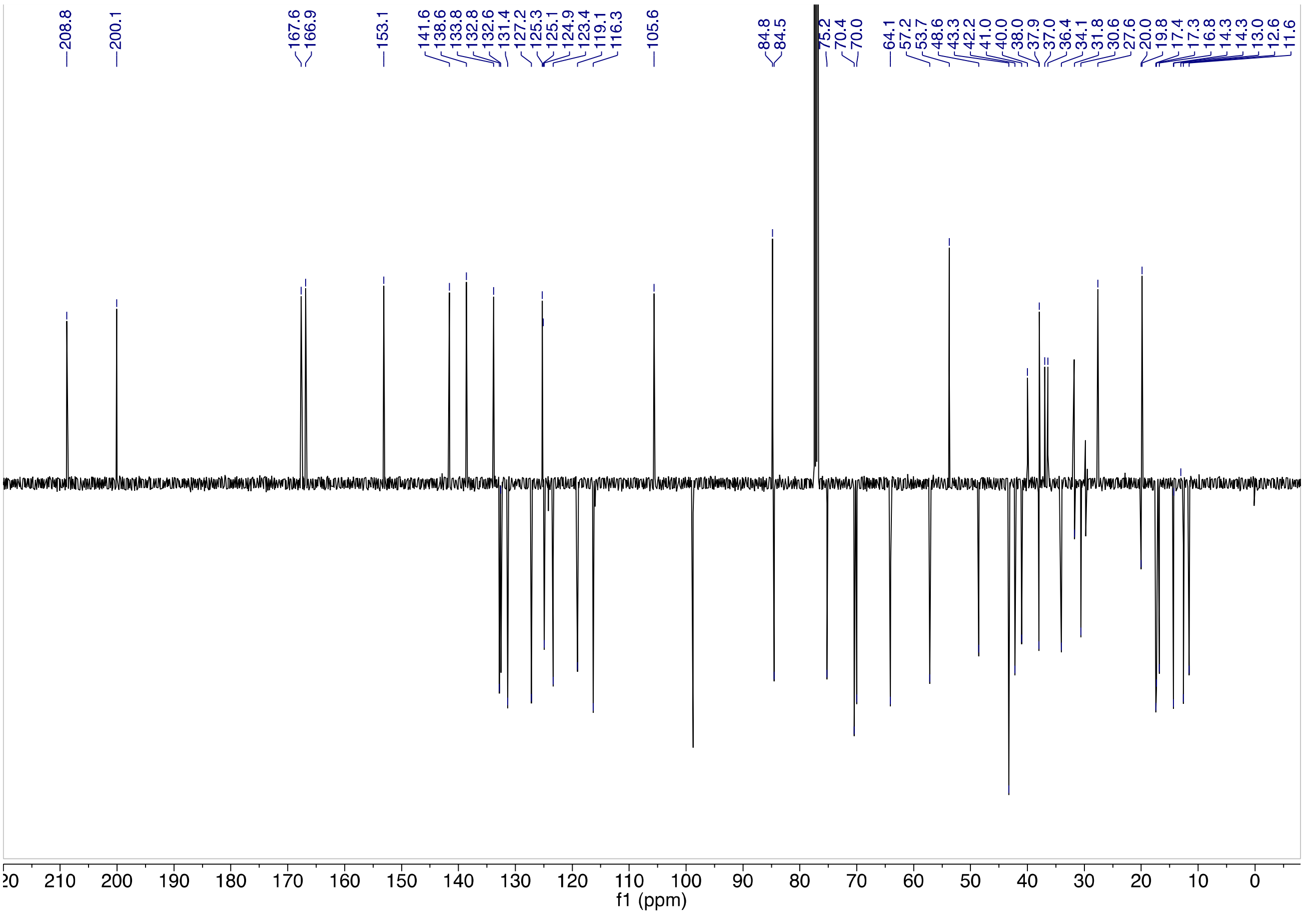

**Figure S5. ¹³C (JMOD) NMR spectrum (125 MHz) of pellemicin.**

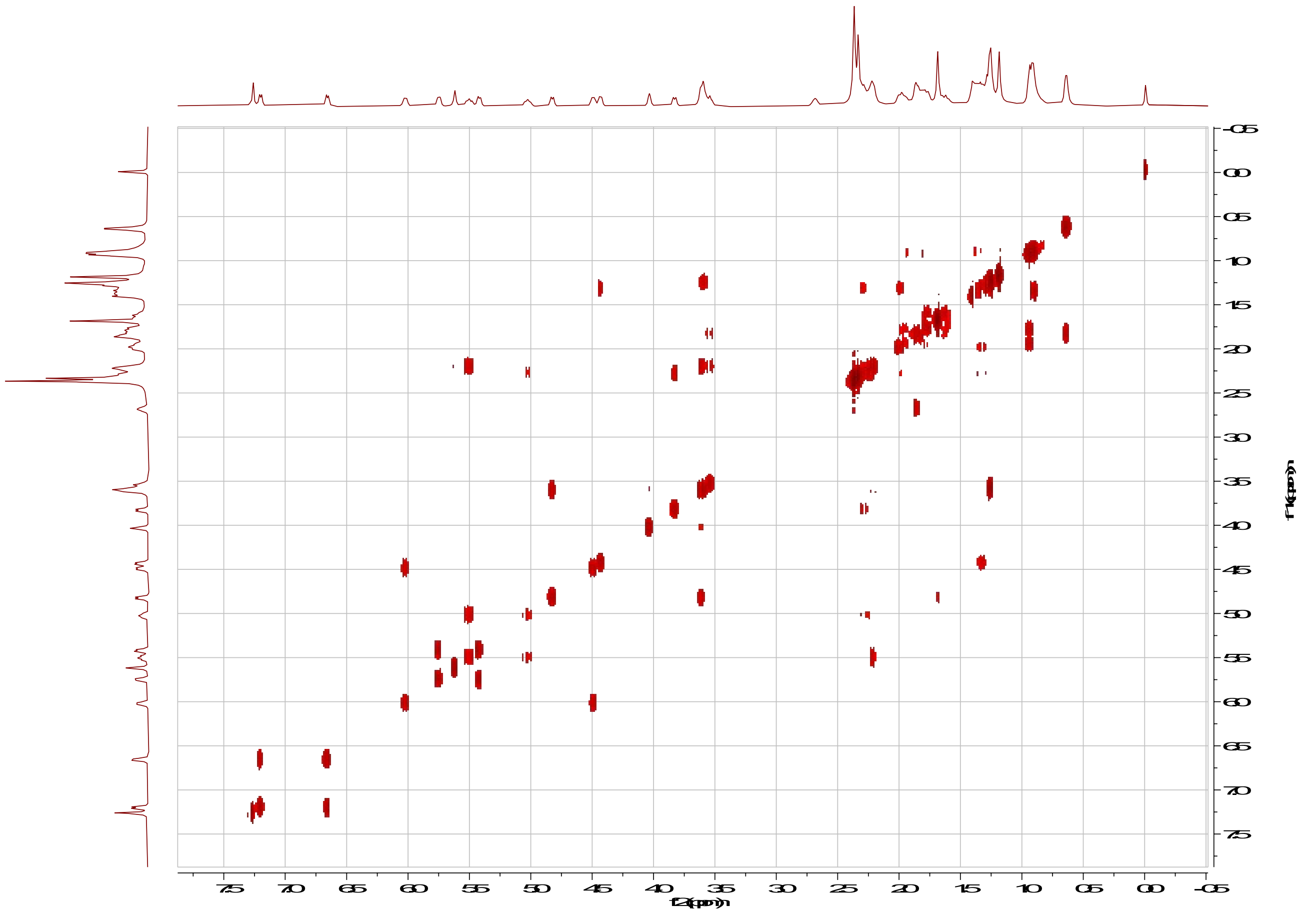

**Figure S6. COSY NMR spectrum (500 MHz) of pellemicin.**

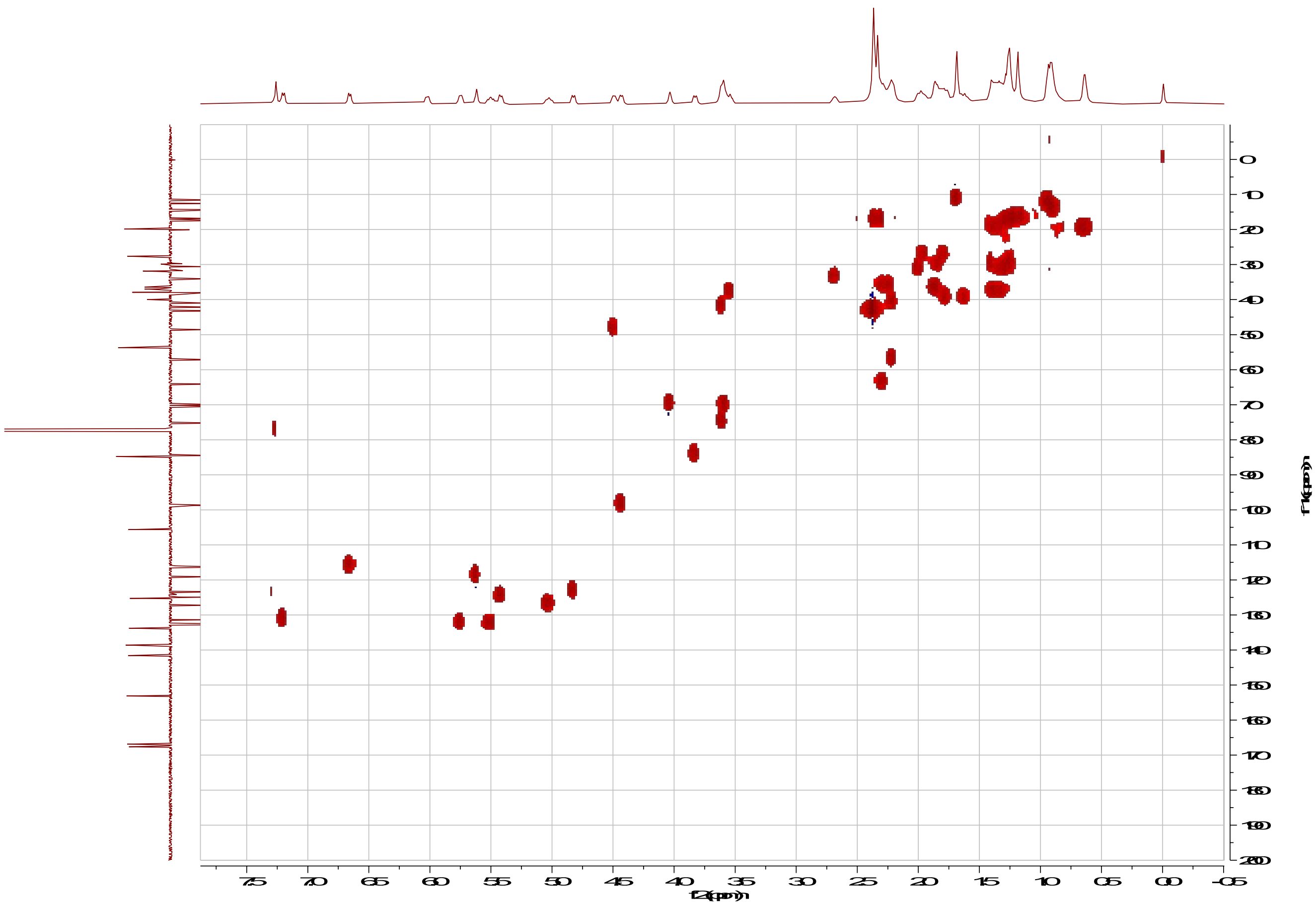

**Figure S7. HSQC NMR spectrum (500 MHz) of pellemicin.**

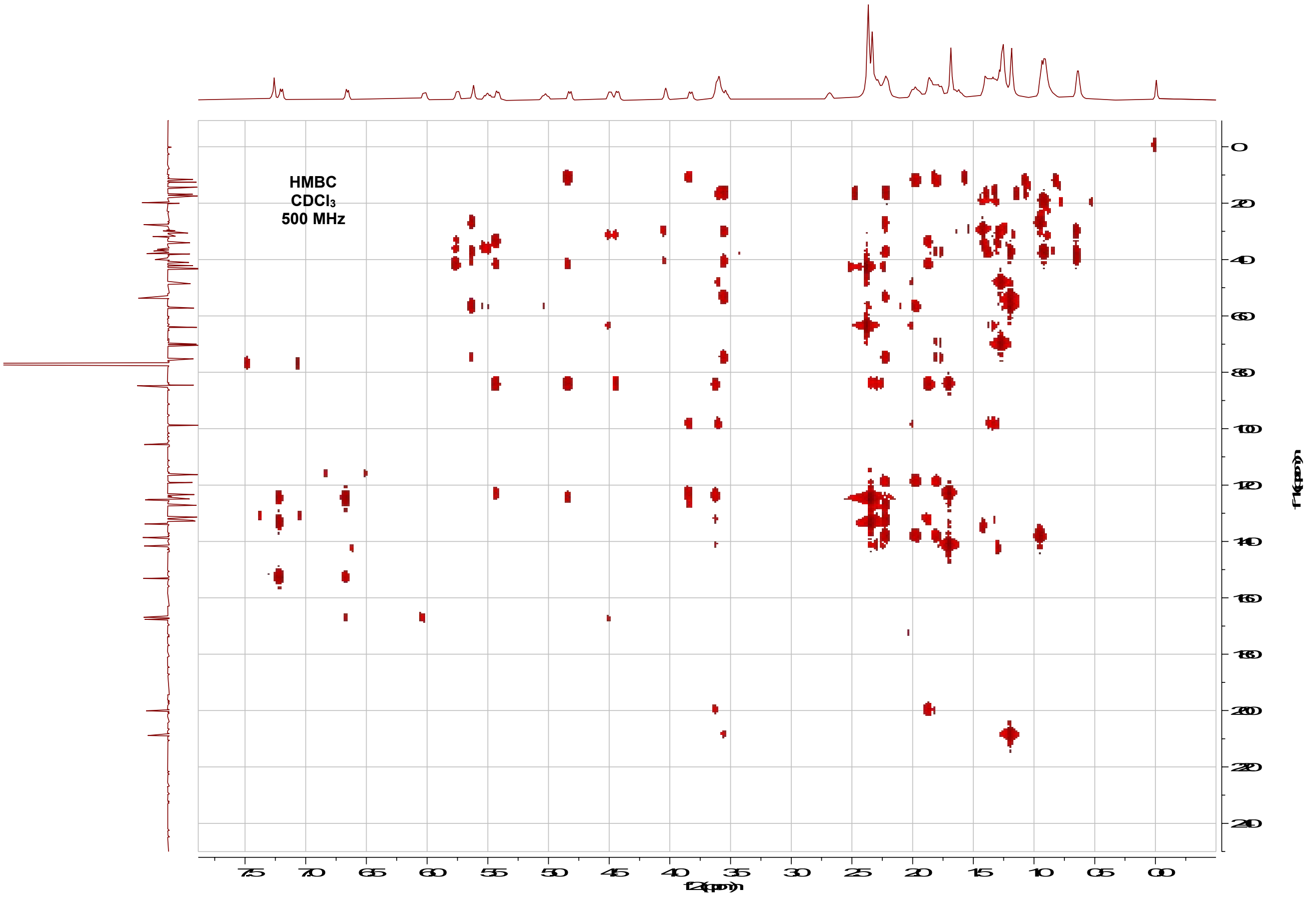

Figure S8. HMBC NMR spectrum (500 MHz) of pellemicin.

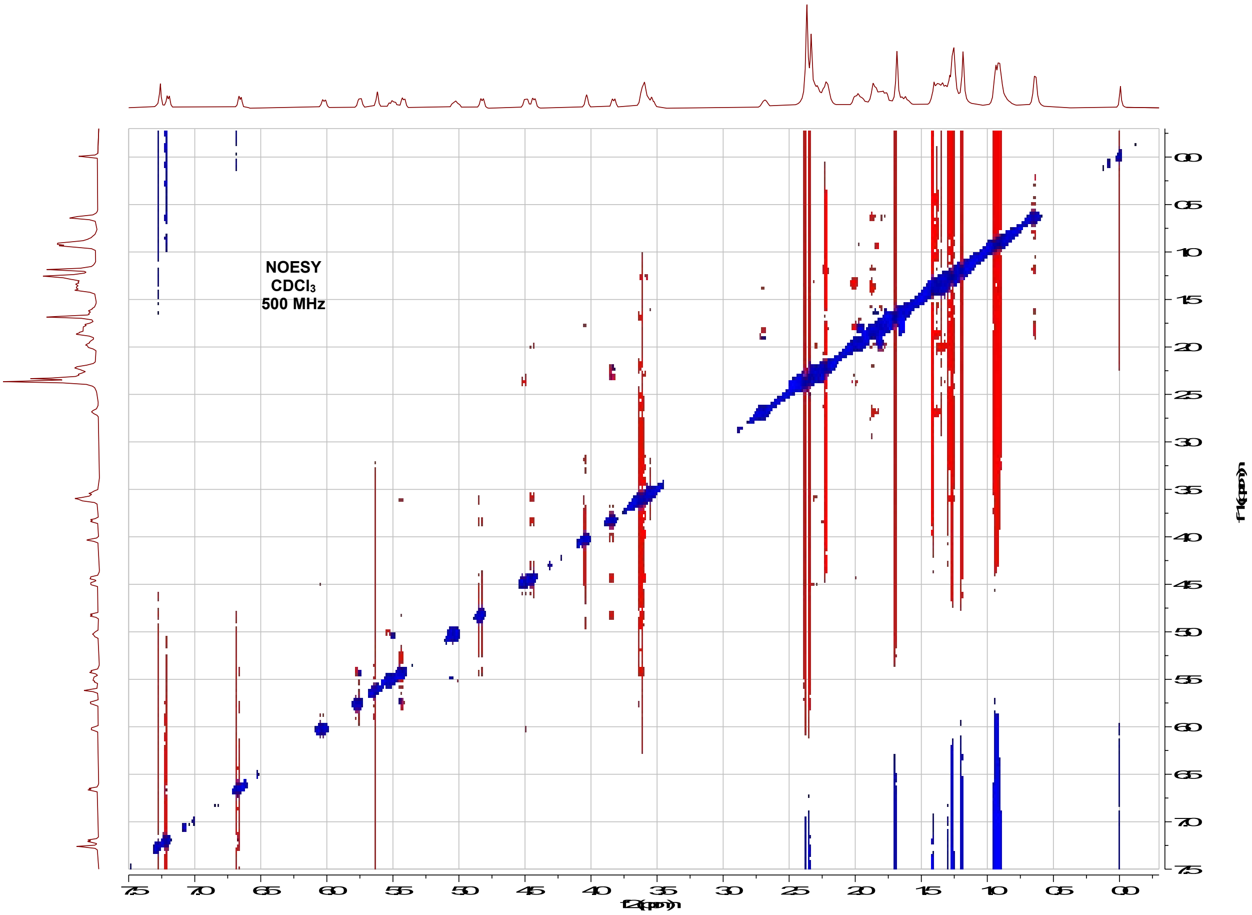

Figure S9. NOESY NMR spectrum (500 MHz) of pellemicin.

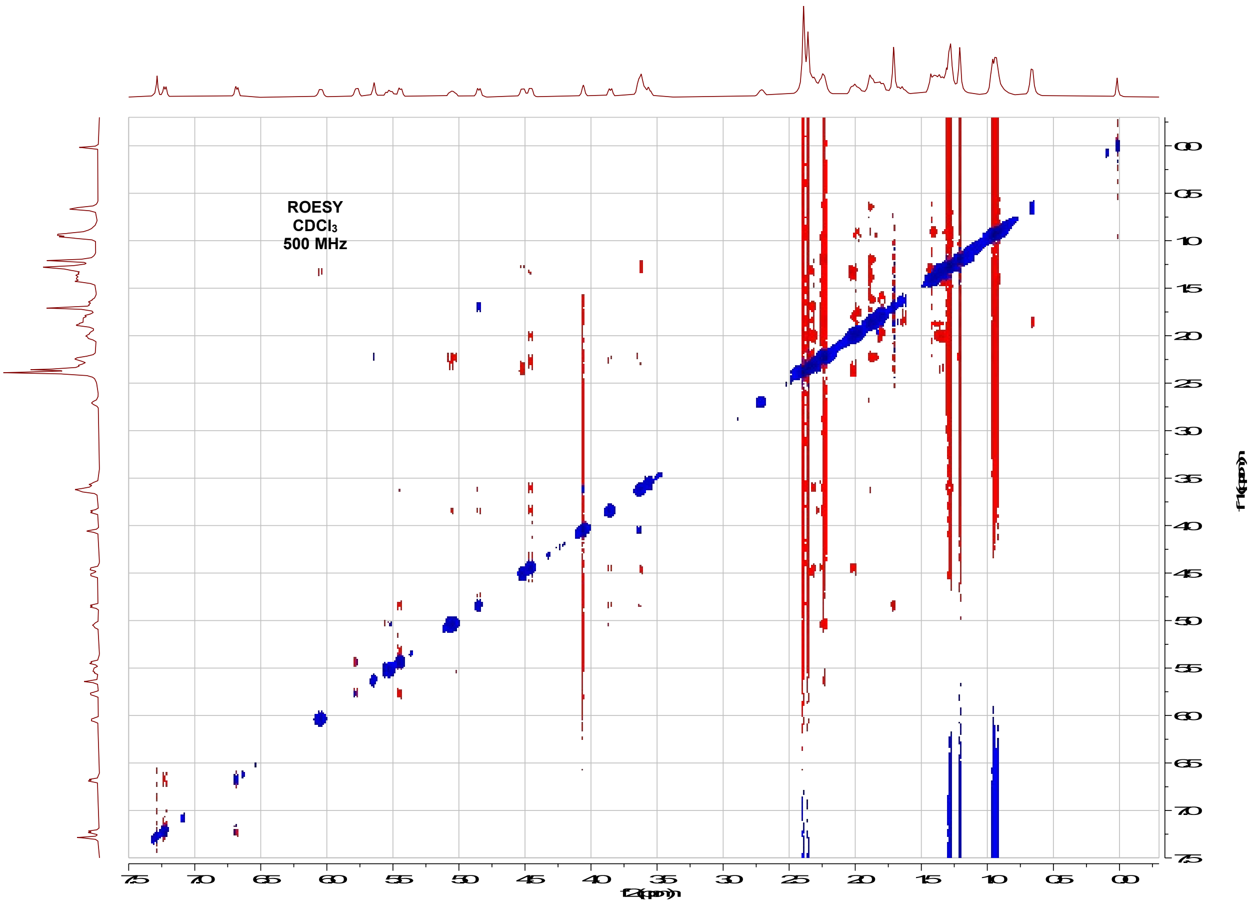

**Figure S10. ROESY NMR spectrum (500 MHz) of pellemicin.**

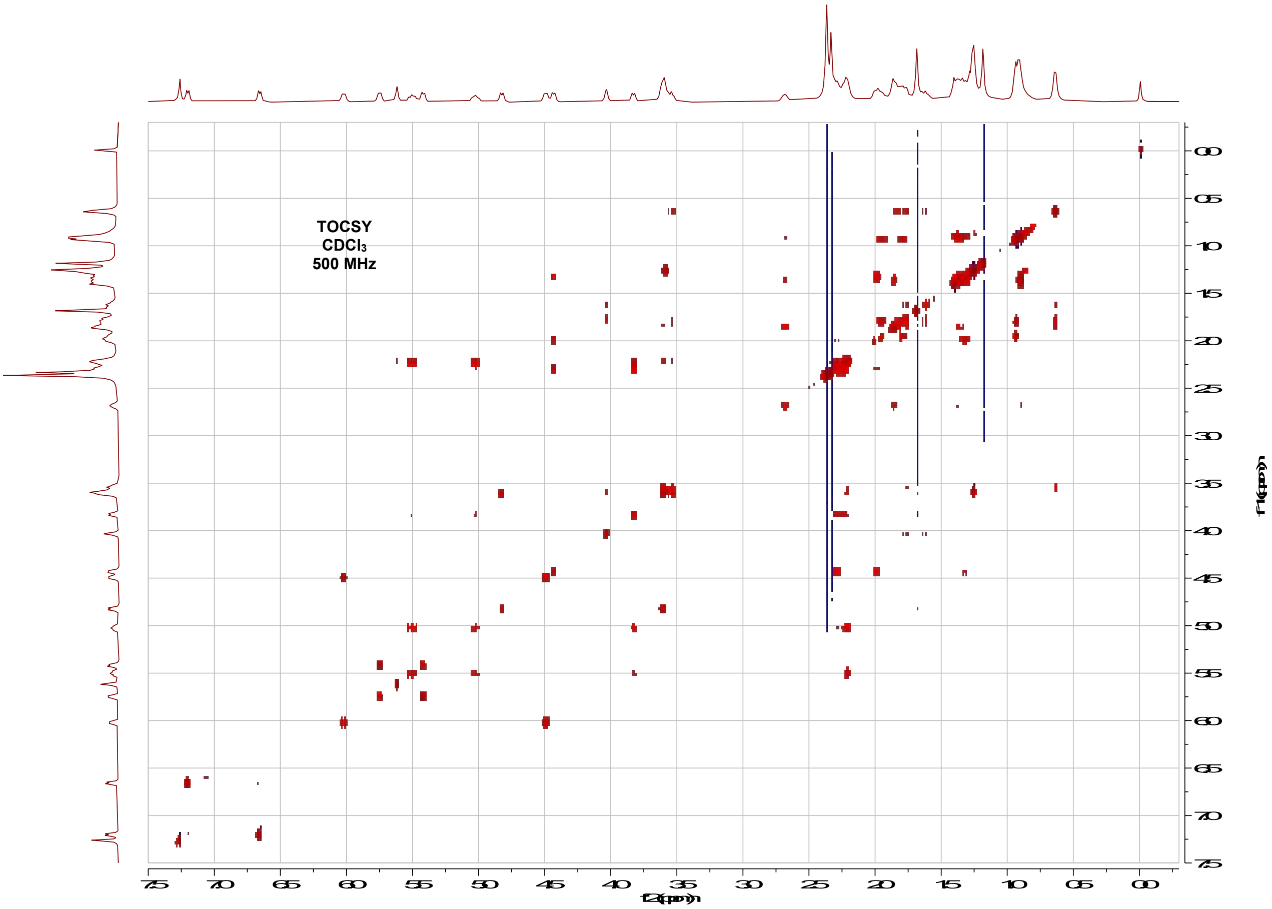

**Figure S11. TOCSY NMR spectrum (500 MHz) of pellemicin**

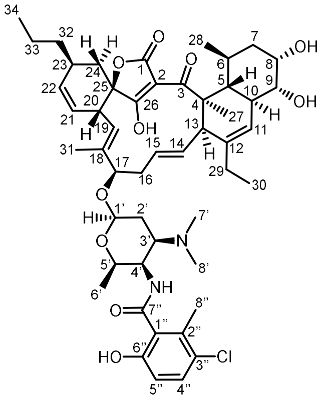

**pellemicin**
literature numbering

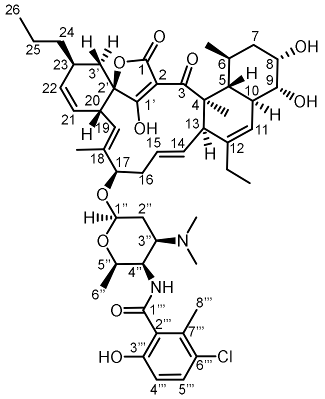

**pellemicin**
biosynthesis-based numbering

**Table S2**: Comparison of $^1$H and $^{13}$C NMR signals for pellemicin isolated in this work (recorded in $CDCl_3$ at 500 / 125 MHz) with those reported by Luk et al. (recorded in $CDCl_3$ at 400 / 100 MHz).[1] Carbons are numbered according to the previous report (left),[1] and the updated biosynthesis-based nomenclature used in Figures 1 and 2 (right).

| Literature C No. | $\delta_C$ / ppm literature[1] | $\delta_C$ / ppm this work | $\delta_H$ / ppm literature[1] | $\delta_H$ / ppm this work | Biosynthesis based C No. |
|---|---|---|---|---|---|
| **1** | 166.8 | 166.9 | | | **1** |
| **2** | 105.6 | 105.6 | | | **2** |
| **3** | 208.7 | 208.8 | | | **3** |
| **4** | 53.7 | 53.7 | | | **4** |
| **5** | 38.0 | 38.0 | 3.54 | 3.55 | **5** |
| **6** | 30.5 | 30.6 | 1.84 | 1.84 | **6** |
| **7** | 39.9 | 40.0 | 1.61 (ax)<br>1.77 (eq) | 1.62 (ax)<br>1.77 (eq) | **7** |
| **8** | 69.9 | 70.0 | 4.02 | 4.03 | **8** |
| **9** | 75.1 | 75.2 | 3.58 | 3.60 | **9** |
| **10** | 40.9 | 41.0 | 2.20 | 2.21 | **10** |
| **11** | 119.1 | 119.1 | 5.63 | 5.62 | **11** |
| **12** | 138.5 | 138.6 | | | **12** |
| **13** | 57.2 | 57.2 | 2.22 | 2.21 | **13** |
| **14** | 132.8 | 132.8 | 5.51 | 5.50 | **14** |
| **15** | 127.1 | 127.2 | 5.02 | 5.02 | **15** |
| **16** | 36.4 | 36.4 | 2.24, 2.32 | 2.24,<br>2.30 | **16** |
| **17** | 84.6 | 84.5 | 3.82 | 3.83 | **17** |
| **18** | 141.6 | 141.6 | | | **18** |
| **19** | 123.2 | 123.4 | 4.82 | 4.82 | **19** |
| **20** | 42.1 | 42.2 | 3.60 | 3.61 | **20** |
| **21** | 124.8 | 124.9 | 5.42 | 5.42 | **21** |
| **22** | 132.4 | 132.6 | 5.75 | 5.75 | **22** |
| **23** | 34.0 | 34.1 | 2.70 | 2.68 | **23** |
| **24** | 36.9 | 37.0 | 1.87 | 1.87 | **3’** |
| **25** | 84.7 | 84.8 | | | **2’** |
| **26** | 199.9 | 200.1 | | | **1’** |
| **27** | 16.7 | 16.9 | 1.19 | 1.18 | **4-Me** |
| **28** | 19.9 | 20.0 | 0.64 | 0.64 | **6-Me** |
| **29** | 27.5 | 27.6 | 1.96, 1.80 | 1.80,<br>1.96 | **12-Et** |
| **30** | 12.5 | 12.6 | 0.94 | 0.94 | **12-Et** |
| **31** | 11.5 | 11.6 | 1.69 | 1.68 | **18-Me** |
| **32** | 37.8 | 37.9 | 1.3-1.4 | 1.32,<br>1.38 | **24** |
| **33** | 19.7 | 19.8 | 1.3-1.4 | 1.34,<br>1.39 | **25** |

| | | | | | |
|---|---|---|---|---|---|
| **34** | 14.2 | 14.4 | 0.91 | 0.91 | **26** |
| **1'** | 98.8 | 98.8 | 4.43 | 4.43 | **1''** |
| **2'** | 31.7 | 31.8 | 2.00 (eq)<br>1.31 (ax) | 2.00 (eq)<br>1.31 (ax) | **2''** |
| **3'** | 64.0 | 64.1 | 2.30 | 2.30 | **3''** |
| **4'** | 48.5 | 48.6 | 4.49 | 4.49 | **4''** |
| **5'** | 70.4 | 70.4 | 3.58 | 3.59 | **5''** |
| **6'** | 17.3 | 17.4 | 1.26 | 1.26 | **6''** |
| **7', 8'** | 43.1 | 43.3 | 2.37 | 2.37 | **3''-NMe$_2$** |
| **4'-NH** | | | 6.04 | 6.02 | **4''-NH** |
| **1''** | 125.2 | 125.3 | | | **2'''** |
| **2''** | 133.7 | 133.8 | | | **7'''** |
| **3''** | 125.0 | 125.1 | | | **6'''** |
| **4''** | 131.2 | 131.4 | 7.21 | 7.20 | **5'''** |
| **5''** | 116.2 | 116.3 | 6.67 | 6.66 | **4'''** |
| **6''** | 153.1 | 153.1 | | | **3'''** |
| **7''** | 167.5 | 167.6 | | | **1'''** |
| **8''** | 17.2 | 17.3 | 2.34 | 2.33 | **8'''** |

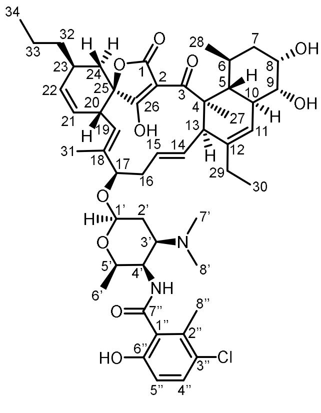

**pellemicin**
literature numbering

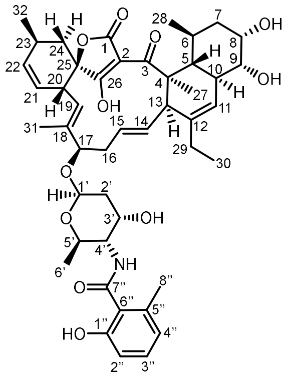

**wychimicin D**
literature numbering

**Table S3**: Comparison of $^{1}H$ and $^{13}C$ NMR signals for pellemicin reported by Luk et al. (recorded in $CDCl_3$ at 400 / 100 MHz) and wychimicin reported by Kimura et al. (recorded in $CDCl_3$ at 600 MHz / 150 MHz).[1,2] Carbons are numbered according to the previous reports.

| Literature C No. | $\delta_C$ / ppm pellemicin[1] | $\delta_C$ / ppm wychimicin D | $\delta_H$ / ppm pellemicin[1] | $\delta_H$ / ppm wychimicin D |
|---|---|---|---|---|
| **1** | 166.8 | 166.8 | | |
| **2** | 105.6 | 105.6 | | |
| **3** | 208.7 | 208.7 | | |
| **4** | 53.7 | 53.6 | | |
| **5** | 38.0 | 37.9 | 3.54 | 3.57 |
| **6** | 30.5 | 30.5 | 1.84 | 1.85 |
| **7** | 39.9 | 39.9 | 1.61, 1.77 | 1.64, 1.79 |
| **8** | 69.9 | 69.9 | 4.02 | 4.05 |
| **9** | 75.1 | 75.2 | 3.58 | 3.63 |
| **10** | 40.9 | 40.9 | 2.20 | 2.23 |
| **11** | 119.1 | 118.9 | 5.63 | 5.63 |
| **12** | 138.5 | 138.7 | | |
| **13** | 57.2 | 57.2 | 2.22 | 2.23 |
| **14** | 132.8 | 132.4 | 5.51 | 5.51 |
| **15** | 127.1 | 127.5 | 5.02 | 5.05 |
| **16** | 36.4 | 36.6 | 2.24, 2.32 | 2.27, 2.33 |
| **17** | 84.6 | 84.4 | 3.82 | 3.84 |
| **18** | 141.6 | 142.1 | | |
| **19** | 123.2 | 122.6 | 4.82 | 4.84 |
| **20** | 42.1 | 41.9 | 3.60 | 3.65 |
| **21** | 124.8 | 124.8 | 5.42 | 5.45 |
| **22** | 132.4 | 133.6 | 5.75 | 5.68 |
| **23** | 34.0 | 29.2 | 2.70 | 2.81 |
| **24** | 36.9 | 38.7 | 1.87 | 1.85, 1.90 |
| **25** | 84.7 | 84.6 | | |
| **26** | 199.9 | 200.0 | | |
| **27** | 16.7 | 16.7 | 1.19 | 1.20 |
| **28** | 19.9 | 19.9 | 0.64 | 0.65 |
| **29** | 27.5 | 27.5 | 1.80, 1.96 | 1.81, 1.98 |
| **30** | 12.5 | 12.5 | 0.94 | 0.95 |
| **31** | 11.5 | 11.5 | 1.69 | 1.74 |
| **32** | 37.8 | 20.9 | 1.3-1.4 | 1.08 |
| **33** | 19.7 | – | 1.3-1.4 | – |
| **34** | 14.2 | – | 0.91 | – |
| **1'** | 98.8 | 96.0 | 4.43 | 4.85 |
| **2'** | 31.7 | 39.3 | 2.00, 1.31 | 1.89, 1.96 |
| **3'** | 64.0 | 67.8 | 2.30 | 4.24 |
| **4'** | 48.5 | 53.2 | 4.49 | 4.06 |
| **5'** | 70.4 | 69.5 | 3.58 | 3.76 |
| **6'** | 17.3 | 18.5 | 1.26 | 1.25 |

| | | | | |
|---|---|---|---|---|
| **7', 8'** | 43.1 | – | 2.37 | – |
| **4'-NH** | – | – | 6.04 | 6.48 |
| **1'' / 6''** | 125.2 | 117.7 | | |
| **2'' / 5''** | 133.7 | 135.2 | | |
| **3'' / 4''** | 125.0 | 122.7 | – | 6.70 |
| **4'' / 3''** | 131.2 | 132.3 | 7.21 | 7.20 |
| **5'' / 2''** | 116.2 | 115.8 | 6.67 | 6.83 |
| **6'' / 1''** | 153.1 | 159.9 | | |
| **7''** | 167.5 | 169.9 | | |
| **8''** | 17.2 | 22.3 | 2.34 | 2.51 |

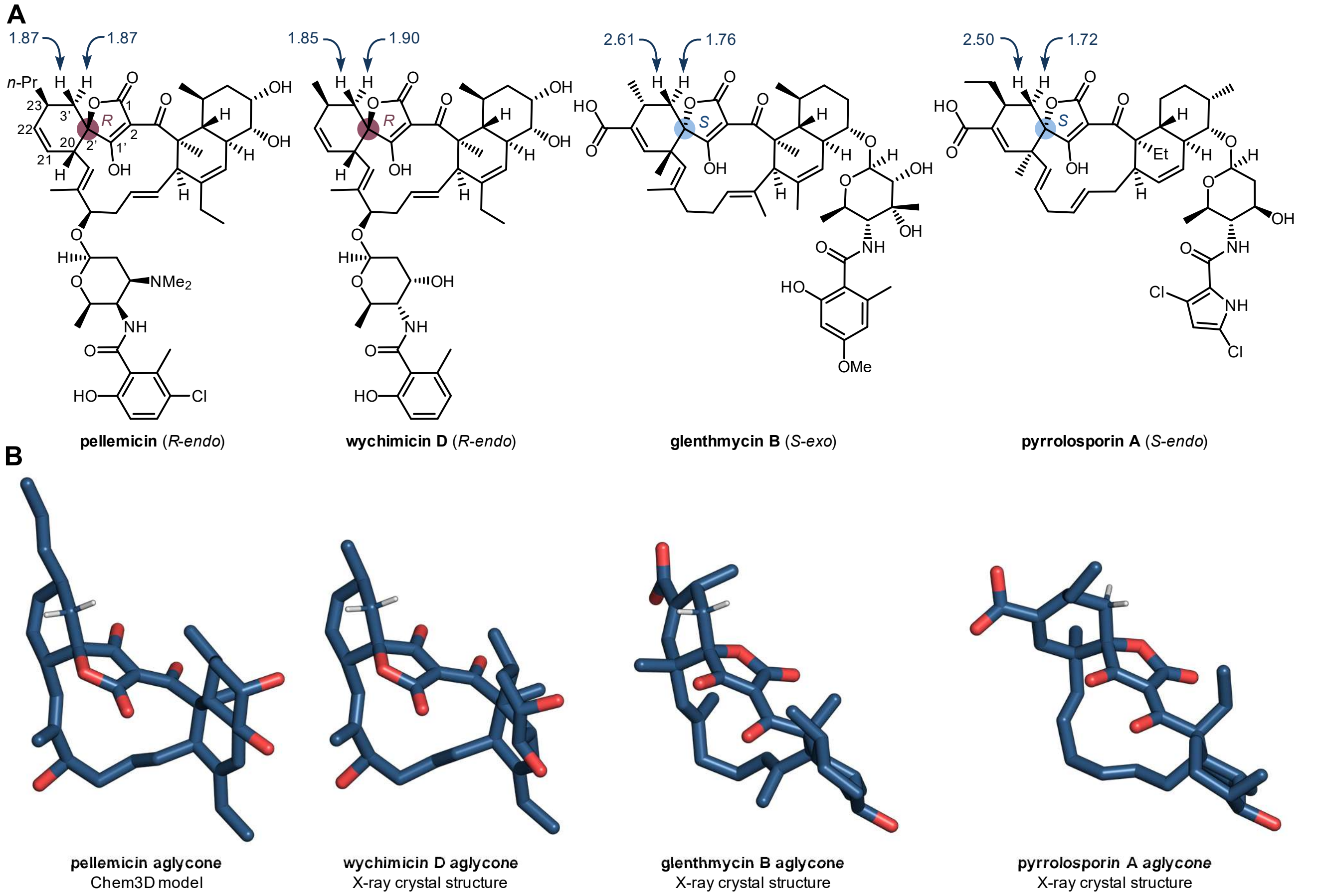

**Figure S12. Comparison of chemical shifts for C24 protons in spirotetronates.** A) Examples of *R-endo* stereochemistry in pellemicin and wychimicin D, *S-exo* stereochemistry in glenthmycin A and *S-endo* stereochemistry in pyrrolosporin A with highlighted $^1$H NMR shift at C24. B) Chem 3D model of pellemicin aglycone and the X-ray crystal structure of wychimicin D,[2] glenthmycin,[3] and pyrrolosporin aglycone.[4] Pyrrolosporin aglycone was manually inverted through the x plane as the original crystal structure was deposited as the incorrect enantiomer.

#### Electronic circular dichroism (ECD) spectra

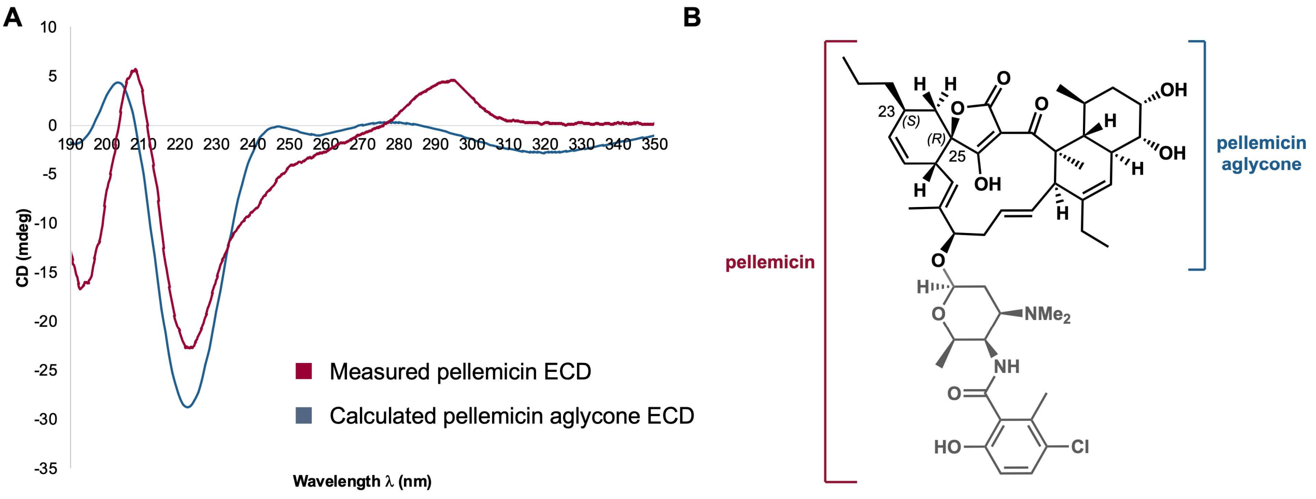

**Figure S13. Electronic Circular Dichroism (ECD) spectra to confirm absolute stereochemistry of pellemicin**. A) Measured Electronic Circular Dichroism (ECD) spectrum of pellemicin and calculated ECD spectrum of (23*S*, 25*R*)-pellemicin aglycone with (*R*)-spiro stereochemistry. B) Structure of (23*S*, 25*R*)-Pellemicin (red bracket) highlighting the aglycone portion in blue brackets.

##### <u>Methods for (23S, 25R)-pellemicin aglycone</u>

##### Method for opt&freq:

%nprocshared=62
%mem=130GB
### opt=tight freq b3lyp/6-311g(d, p) scrf=(iefpcm,solvent=acetonitrile) geom=connectivity empiricaldispersion=gd3bj

##### Method for ECD calculation:

%nprocshared=62
%mem=130GB
### td=(nstates=60) b3lyp/6-311g(d, p) scrf=(iefpcm,solvent=acetonitrile) empiricaldispersion=gd3bj

Optimization & frequence analyses for a total of 27 conformers were accomplished, each of which has the distribution ratio higher than 0.5% (conformers and their distributions are preliminarily generated by Conflex7a using MMFF94 force field); ECD calculation for 20 conformers were accomplished, each of which has the distribution ratio higher than 2.0%

**Table S4. Gibbs free energy and Boltzmann distribution of main conformers of (23*S*, 25*R*)-pellemicin.** To reduce calculation time, conformers with the population over 2% are summarized in the table below and sent to the subsequent ECD calculation.

| Pellemicin | G (Hartree) | Population (%) |
|---|---|---|
| conformer5 | -1849.881436 | 8.563 |
| conformer9 | -1849.881376 | 8.03 |
| conformer15 | -1849.881202 | 6.68 |
| conformer8 | -1849.881177 | 6.504 |
| conformer19 | -1849.88108 | 5.87 |
| conformer25 | -1849.881026 | 5.547 |
| conformer1 | -1849.881021 | 5.515 |
| conformer12 | -1849.880962 | 5.181 |
| conformer2 | -1849.880955 | 5.143 |
| conformer21 | -1849.880935 | 5.035 |
| conformer24 | -1849.880808 | 4.403 |
| conformer4 | -1849.880672 | 3.812 |
| conformer7 | -1849.880589 | 3.492 |
| conformer20 | -1849.880487 | 3.132 |
| conformer13 | -1849.880469 | 3.073 |
| conformer17 | -1849.880464 | 3.058 |
| conformer3 | -1849.880294 | 2.553 |
| conformer26 | -1849.880215 | 2.349 |
| conformer10 | -1849.880161 | 2.219 |
| conformer23 | -1849.880143 | 2.176 |
| conformer6 | -1849.880129 | 2.144 |

**Table S5**: Type II spirotetronate forming Diels-Alderases and the stereochemical course of the reactions they catalyse. Hexacyclospironate is the first biosynthetically-characterised example of an *R*-*endo*-configured type II spirotetronate.[5]

| Compound | Diels-Alderase name | Stereochemical course |
|---|---|---|
| **Decatromicin** | Adec 02955 | *S-endo* |
| **Pellemicin** | BZB76_6464 | *R-endo* |
| **Chlorothricin** | ChlL | *S-exo* |
| **Hexacyclospironate** | HsnL | *R-endo* |
| **Lobophorin** | LobD1 | *S-exo* |
| **Pyrrolosporin A** | PloI4 | *S-endo* |
| **Pyrroindomycin A, B** | PyrI4 | *S-exo* |
| **Tetrocarcins** | TcaU4 | *S-exo* |
| **Versipelostatin** | VstJ | *S-exo* |

#### II. Influenza Assays

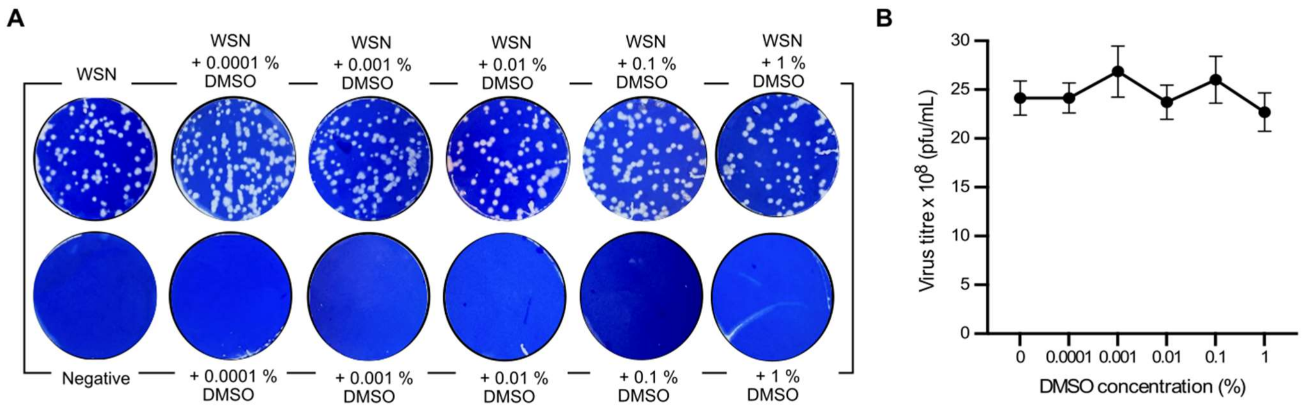

**Figure S14. DMSO does not inhibit influenza induced cell death.** A) MDCK cells were infected with influenza A virus WSN/33 (MOI 0.1) and treated with DMSO (0.0001 – 1%) or left untreated for 1 hour. Cells cultured in DMEM/0.5% FCS alone or DMSO (0.0001 – 1%) without virus were included as controls. Representative images of plaque assays. The cell supernatant was harvested and titred via plaque assay. B) Mean virus titre compared between untreated and treated groups using an unpaired t-test, from three experiments, all samples were non-significant. Error bars show standard deviation.

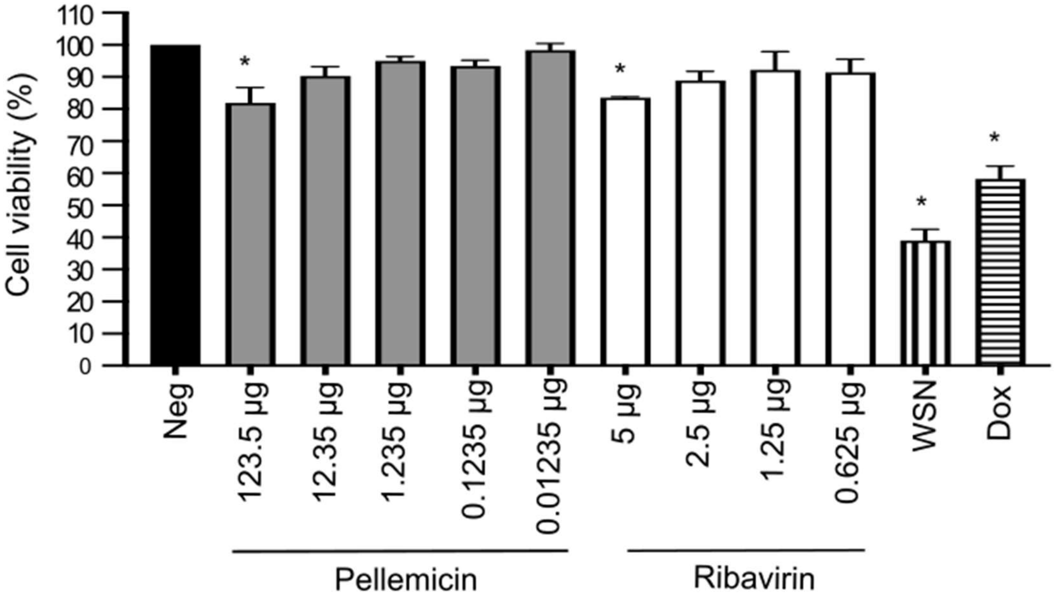

**Figure S15. Cytotoxicity of pellemicin.** MDCK cells were left untreated (negative control), treated with pellemicin (123.5 – 0.01235 µg/mL), ribavirin (5 – 0.625 µg/mL), infected with A/WSN/33 (WSN), or treated with doxorubicin (Dox; 0.04 mM). Cell viability was measured using a neutral red assay kit in triplicate and absorbance measured at 540 nm. The negative control was set at 100% and groups compared with a one-way ANOVA (*p<0.05). Error bars show standard deviation.

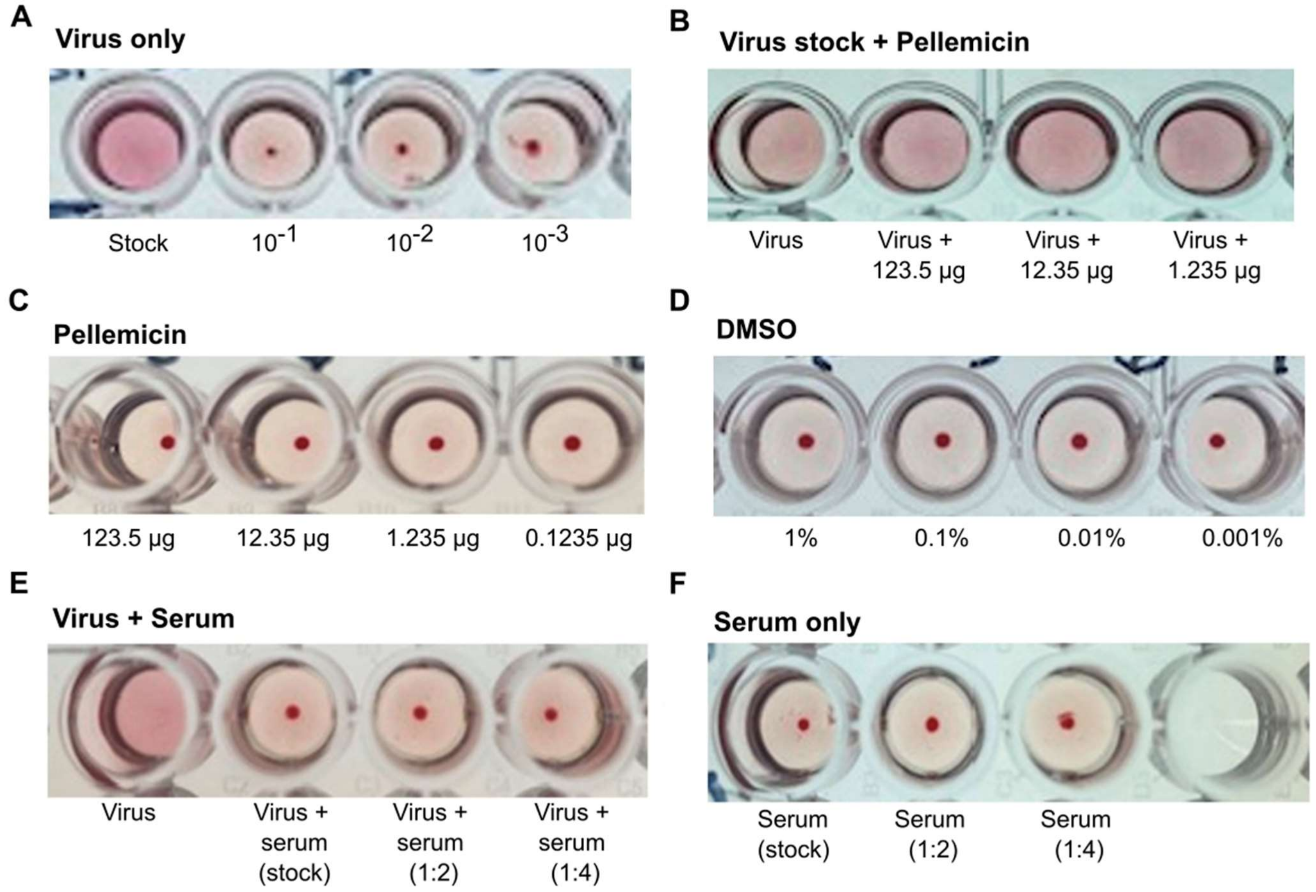

**Figure S16. Effect of pellemicin on agglutination of red blood cells.** A/WSN/33 was mixed with red blood cells in the presence or absence of pellemicin (A&B). Control groups with pellemicin alone (C) or DMSO (D) were included. A positive control with serum from a patient immunised with the influenza vaccine was included, with virus (E) and without (F). Haemagglutination activity was tested by visually assessing haemagglutination.

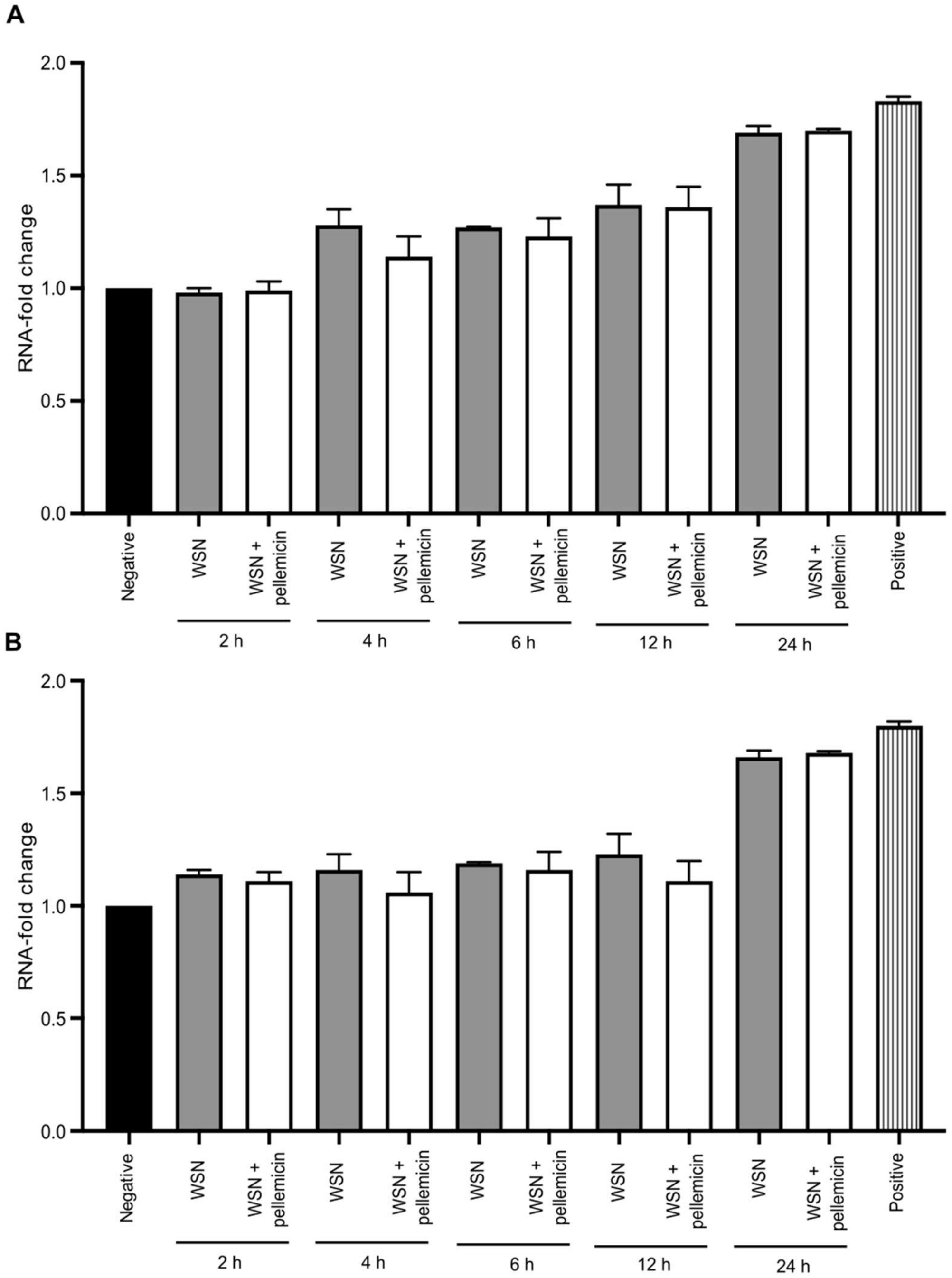

**Figure S17. Pellemicin does not affect viral RNA replication activity.** Quantitative RT-PCR of influenza A cell lysates (A) and supernatants (B). MDCK cells were infected with influenza A/H1N1/WSN (MOI 0.01) and treated with pellemicin (0.1235 µg, white) or left untreated (grey) for 2-24 hours. Nuclease-free water was used as a negative control (black) and a plasmid containing sequences for influenza A as a positive control (striped). Cq values were measured (number of cycles needed to detect target RNA). Target copies were normalised to the negative control and shown as mean fold-change. The untreated groups were compared to treated groups with unpaired t-test ($p<0.05$). Error bars show standard deviation of three independent repeats.

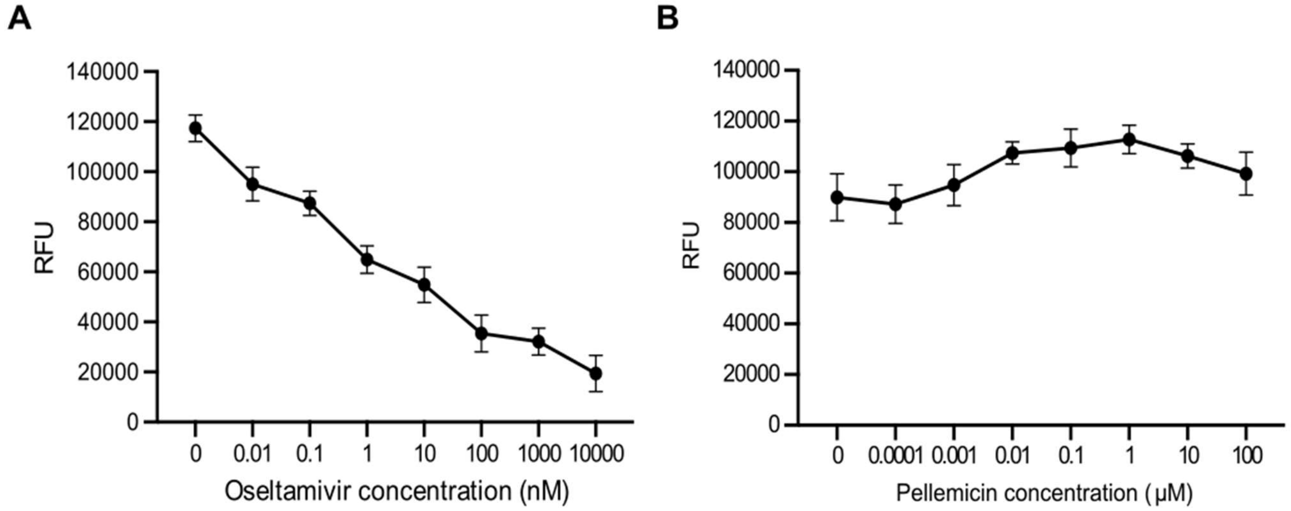

**Figure S18. Pellemicin does not affect neuraminidase activity.** Effect of A) oseltamivir (10000-0.01 nM) and B) pellemicin (100-0.0001 μM) on neuraminidase activity of A/WSN/33 virus expressed as mean relative fluorescence units (RFU) from the production of fluorescent 4-methylumbelliferone. Error bars show standard deviation.
